# Tbx1 ortholog *org-1* is required to establish testis stem cell niche identity in *Drosophila*

**DOI:** 10.1101/2025.05.09.653127

**Authors:** Patrick Hofe, Ariel Harrington, Tynan Gardner, Stephen DiNardo, Lauren Anllo

## Abstract

Stem cells require signals from a cellular microenvironment known as the niche that regulates identity, location, and division of stem cells. Niche cell identity must be properly specified during development to form a tissue capable of functioning in the adult. Here, we show that the Tbx1 ortholog *org1* is expressed in *Drosophila* testis niche cells in response to Slit and FGF signals. *org1* is expressed during niche development and is required to specify niche cell identity. *org1* mutants specified fewer niche cells, and those cells showed disruption of niche-specific markers, including loss of the niche adhesion protein Fas3 and reduced *hedgehog* expression. We found that *org1* expression in somatic gonadal precursors is capable of inducing formation of additional niche cells. Disrupted niche identity in *org1* mutants resulted in niche assembly and functionality defects. We find the conserved transcription factor *islet* is expressed in response to *org1* and show that *islet* functions downstream to mediate niche identity and assembly. This work identifies a novel role for *org1* in niche establishment.

**Summary Statement:** Specification of niche identity is crucial to establish a functional stem cell microenvironment. Using a translatable model, we show that niche expression of *org1* influences *islet* to specify niche identity.

## Introduction

Stem cells play a vital role in maintaining functional tissues. The preservation of stem cells in many tissues is controlled by the stem cell niche, a local cellular microenvironment that supplies signals to maintain stem cell identity, location, and division rates (Li & Xie, 2005). A key concept in stem cell biology is use of diffusible signals to regulate stem cell behaviors. In models like the mammalian intestinal stem cell (ISC) niche, diffusible signals such as Wnt, Hh, and BMP regulate stem cells (Walker et al., 2008). These signals have varying origins since there are multiple cell types that comprise the intestinal stem cell niche (Walker et al., 2008). Because various cell lineages can comprise the niche, identification of niche cells is often challenging. In some tissues even the identity of the stem cells is unclear (Sato et al., 2011; Kaestner, 2019; Li et al., 2023). Thus, identifying regulators that specify niche identity during tissue development has remained a challenge in stem cell biology.

Because of its readily identifiable, well-defined niche, we leverage the *Drosophila* testis to study how niche cells acquire their identity and adopt proper function during development (de Cuevas & Matunis, 2011;Le Bras & Van Doren, 2006;Anllo et al., 2019;Okegbe & DiNardo, 2011;Wingert & DiNardo, 2015). The *Drosophila* testis is a paradigmatic niche in which many concepts at the foundation of stem cell biology continue to be elucidated (Herrera & Bach, 2019; Leatherman & DiNardo 2008 and 2010; Lenhart & DiNardo 2015; Inaba et al., 2015; Tulina & Matunis 2001; Yamashita et al., 2003; Zheng et. al., 2011). The singular cell type that comprises the testis niche, somatic “hub” cells, regulates germline and somatic stem cells required to maintain sperm production throughout the adult life of the fly (Lenhart and DiNardo, 2015; Roach and Lenhart, 2024). The wealth of powerful genetic tools available to study this niche make it an ideal model to identify regulators of niche cell fate.

Prior to niche formation, the embryonic Stage 15 *Drosophila* testis is a spherical arrangement of somatic gonadal precursors (SGPs) interspersed among and encysting germ cells (Jenkins et al., 2003; Le Bras & Van Doren, 2006). SGPs are specified from a subset of mesodermal cells adjacent to muscle precursors in embryonic parasegments 10-13 earlier in embryogenesis (Boyle et al., 1995; Boyle et al., 1997; Riechmann et al., 1998). Niche, or “hub,” cells derive from SGPs, and are initially specified through Notch signaling (Kitadate & Kobayashi, 2010; Okegbe & DiNardo, 2011; DiNardo et al., 2011). By stage 11 of embryogenesis, SGPs contacting the primordial midgut receive Delta ligands which initiate Notch signaling and niche specification (Okegbe & DiNardo, 2011). Continued specification, or further induction of niche cell identity after the initial acquisition of niche cell fate, is required to establish the niche. As part of continued specification, SGPs at the gonad anterior receive the Notch ligand Serrate from other gonadal cells, and SGPs at the posterior receive EGFR antagonistic signals that repress niche identity (Kitadate & Kobayashi, 2010). Defects in Notch signaling result in fewer niche cells specified (Okegbe & DiNardo, 2011; DiNardo et al., 2011; Kitadate & Kobayashi, 2010; Wingert & DiNardo, 2015). Specified niche cells downregulate the conserved Maf transcription factor Traffic jam (Tj) in response to Notch (Wingert & DiNardo, 2015). A role for Notch signaling in niche formation has translated to other niches in *Drosophila* and in mammals (Zamfirescu et al., 2022). Outside of Notch signaling, there is little known about how testis niche cells acquire their identity.

After the initiation of specification, embryonic niche cells are compartmentalized to the gonad anterior in a process called niche assembly (Le Bras & Van Doren, 2006; Anllo et al., 2019). During assembly, specified prospective (pro) niche SGPs move peripherally onto the gonadal extracellular matrix (ECM), where they then migrate to their anterior assembly position (Anllo et al., 2019). Our previous work revealed that extrinsic Slit and FGF signals from neighboring visceral muscle (Vm) are required for this migration (Anllo & DiNardo, 2022). At the gonad anterior, niche cells first exhibit F-actin polarization to niche-niche interfaces, which later re-polarizes to niche-stem cell interfaces as niche cells compact into a spherical arrangement with smooth contour (Anllo & DiNardo, 2022; Warder et al., 2024). Germline stem cells contacting the assembled niche contribute to niche morphogenesis through actomyosin networks that result in tension around the compacting hub periphery (Warder et al., 2024). The assembled and compacted niche expresses niche specific markers such as *fasciclin3* and *hedgehog* and is radially surrounded by germline stem cells (GSCs) and later, cyst stem cells (CySCs) (Le Bras & Van Doren, 2006; Sheng et al., 2009; Sinden et al., 2012; Wingert & DiNardo, 2015). The niche makes adhesive connection to neighboring stem cells and extracellular matrix to maintain its shape, location, and function (Lee et al., 2008; Vida et al., 2024; Tanentzapf et al., 2007; Tseng et al., 2022; Amoyel et al., 2013).

After niche establishment, the niche functions throughout the life of the animal. The niche’s location, critical to maintain GSCs and CySCs, is maintained through Integrin-mediated connections to the ECM (Anllo et al. 2019; Lee et al., 2008; Tanentzapf et al., 2007). In its established position, the niche sends maintenance signals to its surrounding stem cells, including Unpaired (Upd) and Hedgehog (Hh). Upd ligands activate JAK/STAT signaling in GSCs, which regulates their adhesion to the niche and oriented cell divisions (Chen et. al., 2018; Leatherman & DiNardo 2008 and 2010; Tulina & Matunis, 2001; Wawersik et al., 2005; Yamashita et al., 2003). Niche-secreted Hedgehog regulates CySC self-renewal (Amoyel et al., 2012). Niche signals are important for establishing the stem cell populations, and for long term stem cell maintenance (Sheng et al., 2009; Sinden et al., 2012).

Because Notch signaling required to specify niche cells is initiated at stage 11 of embryogenesis (Okegbe & DiNardo, 2011), and niche-specific markers such as Fas3 accumulation are not observed until stage 15 of embryogenesis (Le Bras & Van Doren, 2006; Anllo et al., 2019), there are likely other factors influencing the specification of niche cells during this period. Our work here identifies the Tbx1 ortholog *org1* as one of those factors. Tbx1 and its orthologs play a conserved role across phyla in specifying mesodermal cell types (Marcellini et al., 2003). Like Tbx1*, org1* encodes a transcription factor with a T-box DNA binding domain and is a known regulator of mesodermal muscle fate during embryonic development (Schaub et al. 2012; Schaub & Frasch, 2013; Boukhatmi et al., 2014). Our previous work identified that the conserved transcription factor *islet* is expressed in niche cells and required for both niche assembly and cytoskeletal polarization downstream of Vm assembly cues (Anllo & DiNardo, 2022). *islet* expression is known to be regulated by *org1* during specification of alary musculature (Boukhatmi et al., 2014). Here, we find that a parallel mechanism regulates continued specification of niche cells and directs their anterior assembly. We show that *org1* is expressed in the embryonic niche and reveal its requirement in establishing a full complement of functional, assembled niche cells. We further show that Org1 accumulation is affected by Slit and FGF signals, and that *islet* acts downstream of *org1*. These data reveal *org1* as a novel regulator of niche identity and assembly for the *Drosophila* testis niche.

## Results

### *org1* is expressed in the niche and is induced by Slit and FGF signals

Our previous work has shown that the conserved transcription factor *islet* is expressed in the developing niche (Anllo & DiNardo, 2022). Because Org1 is a known regulator of *islet* in other tissues (Boukhatmi et al., 2014), and single nuclear RNA sequence data (snRNAseq) suggested that *org1* was enriched in the adult testis niche (Li et al., 2021; Raz et al. 2023), we tested whether *org1* was expressed in the developing gonadal niche. Using an Org1::GFP transgenic BAC reporter that rescues *org1* lethality (Kudron et al., 2018 and data not shown), we observed Org1 in prospective niche cells beginning prior to assembly at embryonic stage 15/16 and after niche assembly in stage 17 at the gonad anterior (Figure 1A-1C’). Org1::GFP detection in some prospective niche cells coincided with the timing of Traffic jam downregulation (Figure 1A’, 1B’). Additionally, we also saw SGPs without downregulated Traffic jam that exhibited lower levels of Org1::GFP elsewhere in the gonad (Figure 1B-B”, yellow arrows). We confirmed the stage 17 Org1::GFP expression pattern using an Org1 antibody (Schaub et al., 2012) that revealed a signal in gonadal niche cell nuclei (Figure 1D, D’) which was absent in *org1* mutants (Figure S1). Our findings reveal that *org1* is expressed in the gonad niche, and its expression begins before niche assembly.

**Figure 1.**
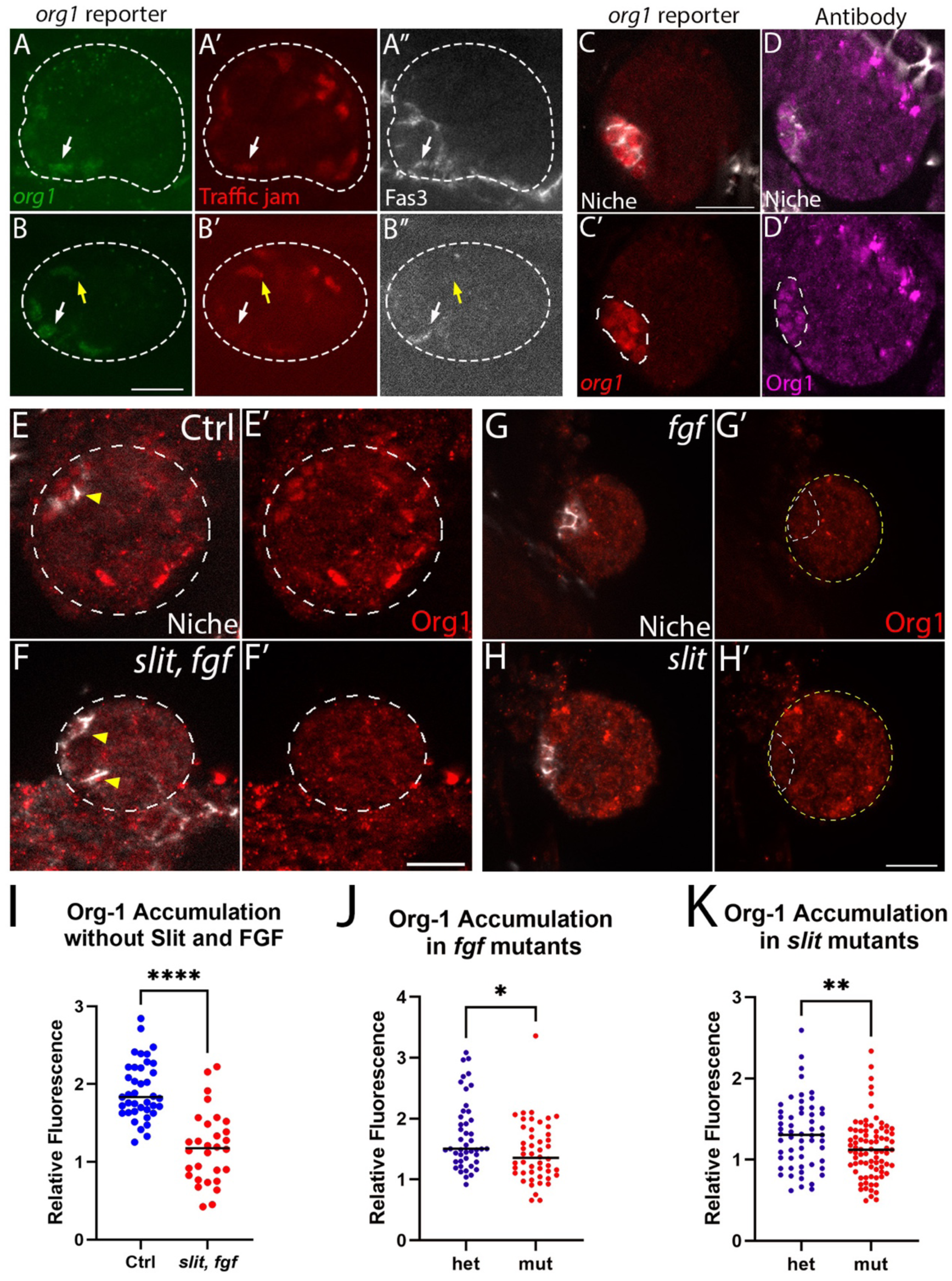
*org1* is expressed in the niche in response to Slit and FGF signals. (A-B’) Gonads (dotted line) from Stage 15 animals with the Org1::GFP BAC reporter, immunostained for GFP (green) and Traffic jam (red). Fas3 (white) accumulates at assembling niche cell interfaces. Org1::GFP is brightest when Traffic jam (red) is downregulated. Prospective niche cells with downregulated Traffic jam (white arrows) and with Traffic jam not downregulated (yellow arrows). (C-H) Gonads dissected from Stage 17 animals aged 20-22 hours at 25 degrees C. (C, C’) Org1::GFP reporter (red) showed *org1* expression in the assembled niche (dotted white line; Fas3, white). (D, D’) Org1 antibody (magenta) revealed accumulation in the niche (dotted white line; Fas3, white) with spurious non-specific signal at the posterior (see Suppl Fig S1). (E-H) Dissected Stage 17 gonads immunostained to reveal Org1 (red) and the Niche (Fas3, white). (E, E’) Control gonads showed Org1 (red) accumulation in the niche (white). (F, F’) *slit, fgf* mutants had diminished accumulation of Org1. Gonads are outlined with dotted lines. (G-G’) *fgf* mutants had diminished niche Org1 accumulation. (H, H’) *slit* mutants had diminished niche Org1 accumulation. Gonads are outlined in yellow; niches are outlined in white. (I) Quantification of Org-1 accumulation in niche cells in controls and Slit and FGF signaling mutants (p<0.0001, Ctrl n=39 niche cells from 13 gonads; *slit, fgf* n= 30 niche cells, from 10 gonads Mann-Whitney Test). (J) Quantified Org1 accumulation in *fgf* mutants and heterozygous sibling controls (Het) (p= 0.0136, Het n=45 niche cells from 15 gonads; *fgf* n=48 niche cells from 16 gonads, Mann-Whitney Test). (K) Quantified Org1 accumulation in *slit* mutants and heterozygous sibling controls (Het) (p= 0.0090, Het n=57 niche cells from 19 gonads; *slit* n=81 niche cells from 27 gonads, Mann-Whitney Test). All quantifications are shown for individual niche nuclei. Scale bars, 10um. n≥3 trials.

Since *islet* is regulated by Slit and FGF signals from the visceral muscle (Vm), we tested if *org1* is also regulated by these signals (Anllo and DiNardo, 2022). If *org1* is induced by these signals, then loss of Slit and FGF Vm signals required for niche assembly would result in loss of Org1 accumulation in niche cells. We assessed Org1 accumulation in double mutant animals with the *slit[2]* mutation and the small chromosomal deficiency Df(2R)BSC25 that removes both genes encoding FGF Heartless ligands, *pyramis* and *thisbe* (see Methods). Indeed, loss of both signals together resulted in diminished Org1 accumulation (Figure 1E-1F’). Additionally, loss of either Slit or FGF separately resulted in diminished Org1 accumulation (Figure 1G-1H’) and our fluorescence quantifications supported these findings (Figure 1I-1K). Together, our data confirm that *org1* is expressed in the niche and accumulates in response to Slit and FGF.

### *org1* is necessary and sufficient for aspects of niche cell identity

Given the known role of *org1* in embryonic muscle specification, we asked if there were niche identity defects in *org1* mutants (Schaub & Frasch, 2012; Boukhatmi et al., 2014). We hypothesized that if *org1* had a role in specifying niche fate, we would see disruption of identity markers. We assayed for niche identity markers Fasciclin3 (Fas3) accumulation and *hedgehog* (*hh*) expression (Le Bras & Van Doren, 2006; Wingert & DiNardo, 2015) in an *org1* mutant allele lacking the translation initiation codon and T-box domain (Schaub et al., 2012). *org1* mutants did not accumulate the niche specific adhesion protein Fasciclin 3 (Figure 2B-2B’). Additionally, *org1* mutants had lower expression of a *hh-*lacZ reporter in niche cells compared to controls (Figure 2C-2E). Interestingly, *hh* expression is known to be restricted to the adult niche (Amoyel et. al., 2013). We find that while *hh* expression is highest in control embryonic niche cells, we detect low levels of *hh* in some control SGPs outside the niche (Figure S2). Since these niche identity markers were disrupted, we used N-cadherin (N-cad) accumulation to detect niche cells in *org1* mutants. N-cad accumulation is enriched at the cortex of properly assembled anterior niche cells, with low accumulation in other SGPs, affording a reasonable measure of identification of niche cells at the anterior (Le Bras & Van Doren, 2006). When visualizing gonads with N-cad immunofluorescence, we detected fewer N-cad accumulating niche cells in *org1* mutants compared to controls (Figure 2F-2G’). The detection of N-cad in some cells indicates that not all aspects of niche identity are disrupted, suggesting a role for *org1* in continued induction of niche fate after their initial specification.

**Figure 2.**
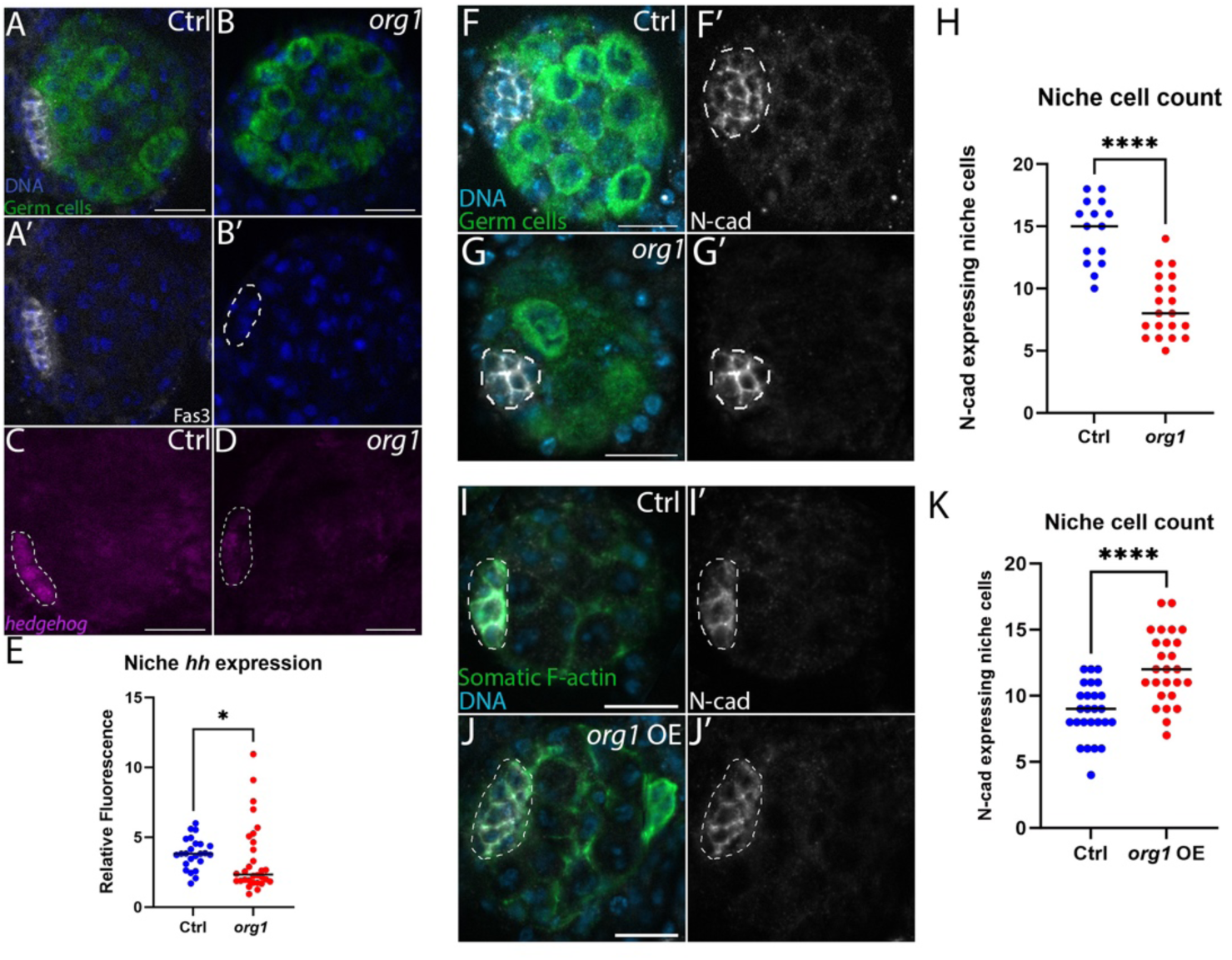
*org1* is necessary and sufficient for aspects of niche cell identity. (A-D) Dissected gonads immunostained to show germ cells (green), Fas3 (white), *hh*-lacZ (magenta), DNA (blue). (A, A’, C) Control niche cells accumulated niche specific markers Fas3 and *hedgehog* reporter expression (*hh*-lacZ, separate gonad from A, A’). (B, B’, D) *org1* mutants did not accumulate Fas3 and had diminished expression of *hh*-lacZ (D is a separate gonad from B). (E) Quantified *hh*-lacZ expression in niche cells accumulating N-cad (p= 0.0294, Ctrl n=24 niche cells from 7 gonads; *org1* n=30 niche cells from 10 gonads, Mann-Whitney Test). (F, F’) Control gonads accumulated enriched N-cad at the cortex of niche cells. (G, G’) *org1* mutant niches had fewer cells accumulating enriched N-cad. (F-G’) Germ cells (green), N-cad (white), DNA (blue). (H) Quantification of N-cad accumulating niche cells per gonad (p<0.0001, Ctrl n=15 gonads; *org1* n=20 gonads, Mann-Whitney Test). (I, I’) Control gonad with *six4*moesinGFP marking somatic F-actin. (J, J’) SGP driven *org1* overexpression in gonads generated niches that had more cells in comparison to controls. (I-J’) Somatic cells (green), Niche (white), DNA (blue). (K) Quantification of niche cell count (p<0.0001, Ctrl n=26 gonads; *org1* n=26 gonads, Mann-Whitney Test). Dotted lines, niche. Scale bars, 10 um. n≥3 trials.

To confirm that the fewer N-cad accumulating niche cells in *org1* mutants resulted from failure of continued niche cell specification in *org1* mutants, and not simply a reduction in the number of SGPs, we quantified the total number of SGPs in *org1* mutants. We assayed the SGP-specific transcription factor Traffic jam (Tj) to count all SGPs (Wingert & DiNardo, 2015). While there was a slight decrease in number of SGPs, the fraction of these cells specified into niche cells was significantly lower in *org1* mutants compared to control gonads (Figure S3). Together with our data indicating a loss of the niche specific adhesion protein Fas3 and reduction of *hh* gene expression in the niche, these results support a role for *org1* in continued specification of niche identity (Figure 2, S3). Interestingly, *org1* mutant niche cells still down regulated *tj* as in normal niches, indicating that some aspects of niche specification occur (Figure S3).

Since *org1* mutants had fewer niche cells, we tested whether overexpression of *org1* in all SGPs would create more niche cells. Indeed, when overexpressing *org1* with a somatic lineage-specific driver (*six4-*gal4; Anllo et. al., 2019), we generated additional N-cad and Islet accumulating niche cells at the gonad anterior (Figure 2I-K; 5E-F). Together, our data suggest that *org1* plays a role in specifying niche identity and is sufficient to induce niche identity in additional SGPs.

### *org1* is required SGP intrinsically for niche identity and assembly

The data above were collected from a whole organism *org1* mutant line in which all T-box binding domains were excised (Schaub et al., 2012). To test whether *org1* is required autonomously in SGPs, we disrupted *org1* using two somatic lineage-specific drivers that varied in strength, *six4*VP16-gal4 (Warder et al., 2024) and *six4*-gal4. Using the weaker driver (*six4-* gal4), we found an expected decrease in niche cell number, and Fas3 accumulation appeared reduced (Figure S4). Assuming this driver generates only a partial knockdown, this suggests a stringent requirement of *org1* in niche cell specification (See Discussion). Excitingly, using the stronger driver (*six4*VP16-gal4), we saw both fewer niche cells (Figure 3A-3C), and dispersed niche cells accumulating low levels of Fas3 (Figure 3B’-3B’’, 3D). It is possible this dispersed phenotype was not something we could observe in the global *org1* mutants due to the complete loss of Fas3 (Figure 2). Other niche cell markers, including E-cad and N-cad, accumulate on control non-niche SGPs to a degree, so dispersed niche cells might not have been apparent (Figure S5). Our data suggest that *org1* is primarily a specification factor during niche establishment and secondarily affects niche assembly.

**Figure 3.**
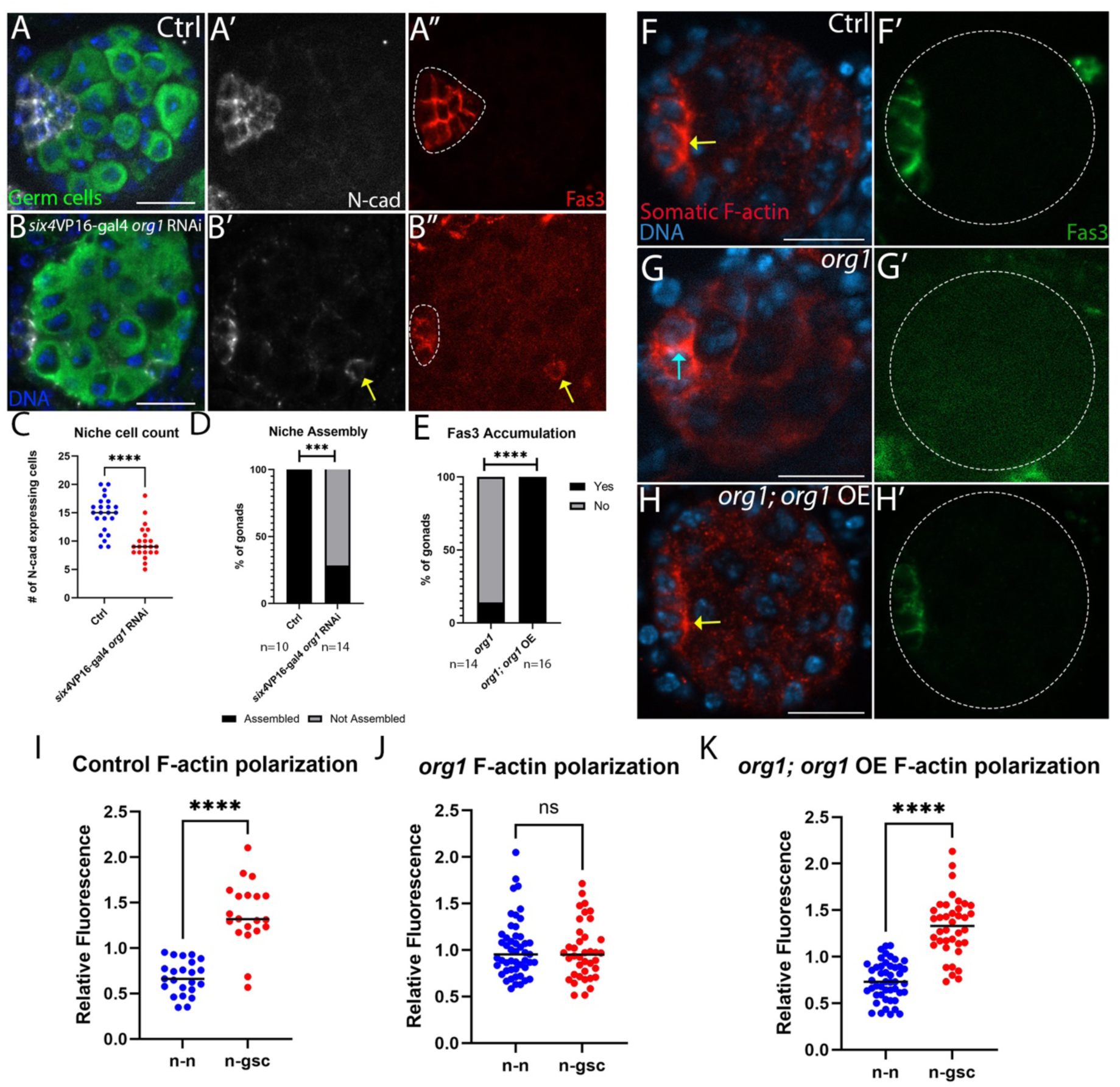
*org1* is required SGP intrinsically for niche identity and assembly. (A, A’, A’’) Control gonads with N-cad (white) and Fas3 (red) accumulating at the anterior. (B, B’, B’’) *org1* knockdown with six4gal4VP16 gonads showed fewer niche cells that accumulate N-cad (white) and Fas3 (red) at the anterior, with displaced niche cells (yellow arrow). Brightness and contrast were adjusted in A” and B” to enable visualization of Fas3. (C) Niche cell count per gonad in *org1* knockdown (p<0.0001, Ctrl n=23 gonads; six4VP16-gal4 *org1* RNAi n=22 gonads, Mann-Whitney Test). (D) Quantification of niche assembly in six4VP16-gal4 *org1* RNAi (p= 0.0006, Ctrl n= 10 gonads; *org1* RNAi n=14 gonads, Fisher’s Exact Test) (E) Quantified Fas3 accumulation in *org1* rescue niches (p<0.0001, *org1* n= 14 gonads; *org1; org1* OE n= 16 gonads, Fisher’s Exact Test). (F-H) Yellow arrows indicate niche-GSC F-actin interfaces and blue arrows indicate niche-niche F-actin interfaces. (F, F’) Control gonad with Fas3 (green) accumulating at the anterior. (G, G’) *org1* knockout mutant gonad with no Fas3 (green) accumulation. (H, H’) *org1* knockout mutant with *org1* overexpression in somatic gonadal cells (red) with rescued Fas3 (green) accumulation. (F-H) arrows indicate F-actin enrichment at specific SGP interfaces. (I) Quantified F-actin polarization in control gonads (p<0.0001, n-n interfaces n=23; n-gsc interfaces n=20, from 7 gonads, Mann-Whitney Test). (J) Quantified F-actin polarization in *org1* mutants (not significant, n-n interfaces n=49; n-gsc interfaces n=38, from 14 gonads, Mann-Whitney Test). (K) Quantified F-actin polarization in *org1* mutant rescued gonads (p<0.0001, n-n interfaces n=47; n-gsc interfaces n=38, from 16 gonads, Mann-Whitney Test). Scale bars, 10 um. n≥3 trials.

To further support an SGP intrinsic requirement for *org1* during niche establishment, we performed a tissue-specific rescue experiment. We expressed *org1* with our somatic lineage specific driver, *six4-*gal4, in whole organism *org1* mutants and observed rescued Fas3 accumulation (Figure 3E-3H’). Our rescue experiment simultaneously expressed the F-actin label Ftractin-tdTomato in SGPs, allowing analysis of niche cytoskeletal polarity. Our previous work has shown that F-actin polarizes to niche-stem cell interfaces after assembly is complete (Anllo & DiNardo, 2022; Warder et al., 2024). While F-actin polarization at niche-stem cell interfaces is disrupted in *org1* mutant niche cells, this polarization was rescued with *org1* overexpression in SGPs (Fig 3F-K, see arrows). These data suggest that *org1* in SGPs is sufficient to initiate downstream cell biological events that polarize F-actin in the assembled niche.

### *org1* is important for niche to stem cell communication

Given the defects in niche identity and organization seen when *org1* is compromised, we tested if the function was compromised without *org1*. We used known steady state niche signaling pathways and centrosome position to assay for niche function deficits (Leatherman & DiNardo, 2010; Kiger et al., 2001; Tulina & Matunis, 2001; Yamashita et al., 2003, 2005). Stat accumulates in GSCs in response to the niche signal Upd1 (Chen et al., 2018; Kiger et al., 2003). We assessed *upd1* expression in niche cells by quantifying levels of *upd1*-Gal4 driven RFP fluorescence and found significantly reduced expression in niche cells in whole organism *org1* mutants (Figure 4A-C). Additionally, we measured reduced Stat accumulation in GSCs (Figure 4D-4E’, 4F), which was consistent with loss of *upd1* expression and impaired niche-GSC communication. Niche cells are also responsible for sending Hh signals to prospective CySCs to enable their self-renewal (Michel et al., 2012; Amoyel et al., 2013). We examined localization of Patched, a Hh receptor that accumulates in vesicles in the niche and CySCs in response to signaling (Amoyel et al., 2013; Michel et al., 2012; Wingert & DiNardo, 2015) (Fig 4G-4H’). *org1* mutants had significantly reduced Patched accumulation in the niche and never accumulated Patched in nearby somatic cells (Figure 4G-4H’, 4I). This finding aligns with the reduced *hh* expression we detected in niche cells when *org1* was disrupted (Figure 2C-E), correlating reduced ligand expression to disruption in niche-CySC communication.

**Figure 4.**
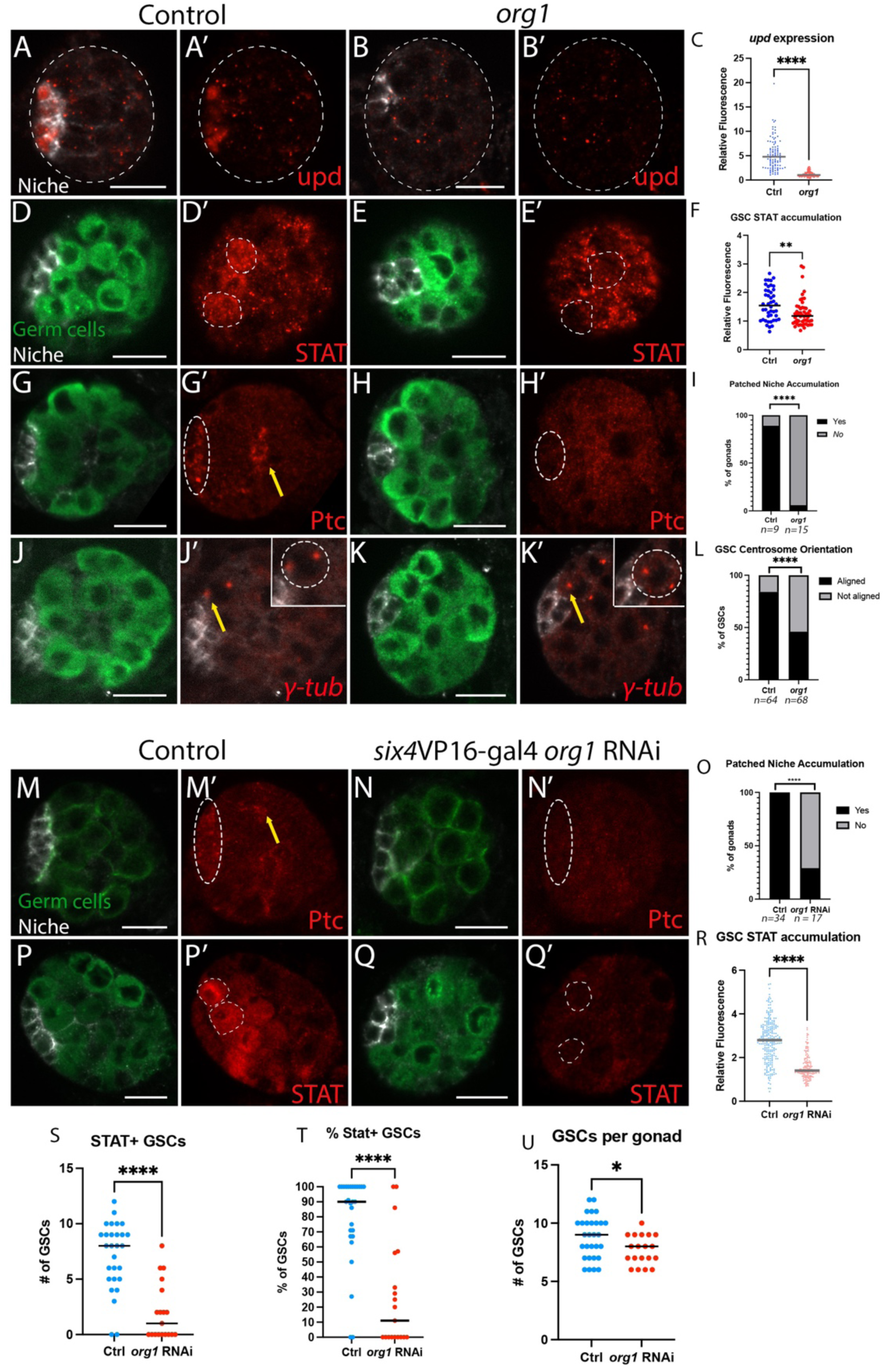
*org1* is important for niche to stem cell communication. (A, A’) *upd1*-Gal4 drives RFP expression in control niche cells, but (B, B’) not in *org1* null animals. (C) Quantification of RFP fluorescence in niche cells (p < 0.0001, Ctrl n = 100 niche cells from 20 gonads; *org1* n = 140 niche cells from 28 gonads) (D, D’) STAT (red) accumulated in germline stem cells (GSCs, dotted line) in control gonads, but (E, E’) not in *org1* mutant germline stem cells. (F) Quantification of STAT accumulation (p= 0.0041, Ctrl n=44 GSCs from 11 gonads; *org1* n=52 GSCs from 16 gonads, Mann-Whitney Test). (G, G’) Hedgehog signaling target Patched (red) accumulated in the niche (white dotted line) and Cyst stem cells (yellow arrow). (H, H’’) Patched did not accumulate in *org1* mutant gonads in the niche or cyst stem cells. (I) Quantified Patched accumulation in the niche (p<0.0001, Ctrl n=9 gonads; *org1* n= 15 gonads, Mann-Whitney Test). (J, J’) In control gonads with duplicated GSC centrosomes, one centrosome was aligned adjacent to the interface with the niche (yellow bracket). Insets show germline stem cells outlined in white, and the niche boundary (Fas3, white). (K, K’) In *org1* mutant gonads, duplicated centrosomes were often not aligned adjacent to the niche interface (yellow bracket). (L) Quantified centrosome alignment (p<0.0001, Ctrl n=64 GSCs from 18 gonads; *org1* n= 68 GSCs from 21 gonads, Fisher’s Exact Test). (J’, K’) Insets show centrosome pairs. (M-N) Hedgehog signaling target Patched accumulated in niche cells and CySCs in controls (M-M’) but not in six4VP16-gal4 *org1* RNAi gonads (N-N’). (O) Quantified Patched accumulation in the niche (p<0.0001, Ctrl n=34 gonads; *org1* RNAi n= 17 gonads, Mann-Whitney Test). (P, P’) STAT (red) accumulated in GSCs (dotted line) in control gonads, but (Q, Q’) not in six4VP16-gal4 *org1* RNAi germline stem cells. (R-T) Quantified STAT accumulation (R), STAT+ GSCs (S), and percentage of GSCs that are STAT+ (T) (p<0.0001, Ctrl n=246 GSCs from 28 gonads; *org1* RNAi n=146 GSCs from 19 gonads, Mann-Whitney Test). (U) Number of GSCs contacting N-cad positive niche cells in control and six4VP16-gal4 *org1* RNAi gonads (p = 0.033, Ctrl n = 29 gonads, six4VP16-gal4 *org1* RNAi n = 19 gonads). For all panels: N-cad (white), scale bars, 10 um. Germ cells are indicated with *nos*-lifeactin::tdTomato (green, panels M-N) or Vasa (green, all other panels). n≥3 trials.

We also tested stem cell behavior in the absence of *org1*. During normal GSC divisions, one centrosome is anchored close to the niche-germline stem cell interface, while the other centrosome moves to the opposite GSC pole. The alignment of one centrosome at the niche interface enables GSC divisions perpendicular to the niche (Yamashita et al., 2003). Gonads from *org1* null animals showed defects in centrosome alignment, with both GSC centrosomes often displaced from the interface with the nearby niche cell. This displacement positioned centrosomes in orientations that would preclude perpendicular GSC divisions from the niche (Figure 4J-4K’, 4L). Together, our data suggest that *org1* is required to generate niche cells capable of communication with stem cells, further supporting a role for *org1* in niche cell specification.

To ask whether *org1* is required cell autonomously in the somatic gonad to enable niche communication with the stem cells, we assessed STAT and Patched accumulation in the six4VP16-gal4 *org1* RNAi condition. Consistent with *org1* null animals, somatic gonadal knockdown with *org1* RNAi resulted in failed accumulation of Patched in the niche and neighboring somatic cells (Figure 4M-O), and failed accumulation of STAT in neighboring GSCs (Figure 4P-R). These data indicate a cell autonomous requirement for *org1* in communication with stem cells.

We hypothesized that failed communication to neighboring stem cells would adversely affect stem cell maintenance. Neither *org1* null nor six4VP16-gal4 *org1* RNAi animals are viable past embryogenesis, precluding analysis of later stem cell maintenance. We thus assessed GSC maintenance by quantifying the number of STAT+ GSCs adherent to the embryonic niche. We determined a threshold level of STAT+ accumulation quantified as one standard deviation below the mean in control GSCs (see methods). Our data revealed significantly fewer STAT+ GSCs (Figure 4S-T), and fewer total GSCs adherent to the niche (Figure 4U) in the six4VP16-gal4 *org1* RNAi somatic cell knockdown, which disrupts *org1* during niche specification and prior to and through niche assembly. These data suggest functional defects in GSC maintenance when *org1* is autonomously disrupted in the somatic gonad.

### *islet* regulates niche cell specification downstream of *org1*

Our previous work has shown that *islet* is expressed downstream of Vm signals Slit and FGF and is important for niche assembly (Anllo & DiNardo, 2022). Here, we show *org1* is also expressed downstream of Slit and FGF and has a role in niche assembly (see Figure 1,3). Because *org1* is known to induce *islet* expression in embryonic muscle (Boukhatmi et al., 2014), we next tested if *org1* regulated *islet* in the embryonic gonad. Indeed, we saw that Islet was no longer enriched in *org1* mutant niche cells (Figure 5A-5C). We also hypothesized that if *org1* is regulating *islet*, we would see Islet accumulation in the additional niche cells specified with our SGP lineage specific *org1* overexpression. Indeed, we saw that additional niche cells at the anterior also accumulated Islet (Figure 5D-5F). These data indicate that Islet accumulates in response to *org1* expression.

**Figure 5.**
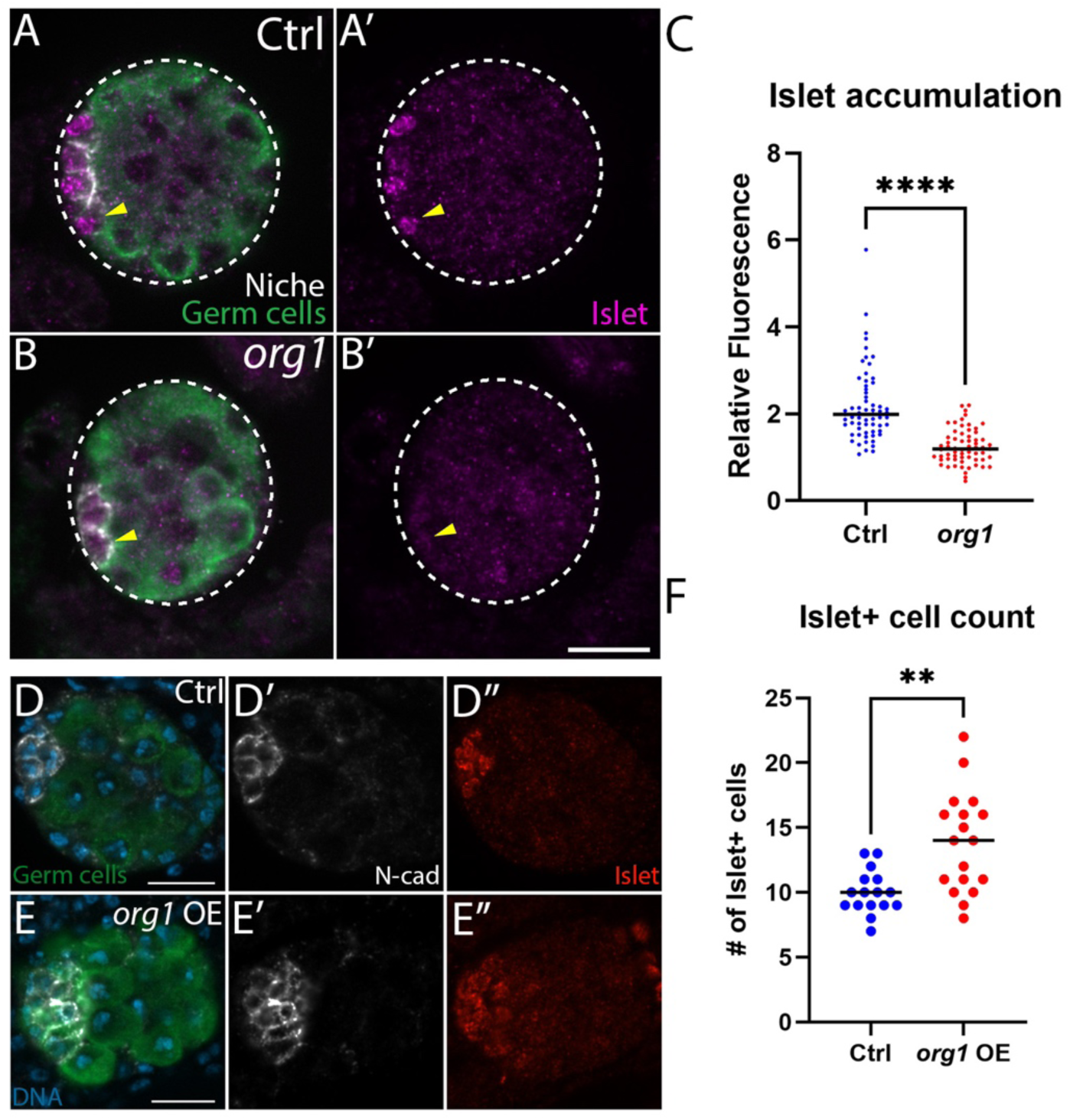
*org1* is important for Islet accumulation. (A, B) Germ cells (Vasa, green), Niche (Ncad, white), DNA (Hoechst, blue). (A, A’) Control gonads showed Islet (magenta) accumulation enriched in the niche. (B, B’) *org1* mutant gonads exhibited diminished Islet accumulation. (C) Islet accumulation quantification in niche cells (p<0.0001, Ctrl n=63 from 21 gonads; *org1* n=63 from 21 gonads, Mann-Whitney Test). (D, D’, D’’) Control gonads with N-cad (white) and Islet (red) accumulating in niche cells. (E, E’, E’’) *org1* overexpressing gonad with additional niche cells (white) that also accumulated Islet (red). (F) Quantification of number of Islet accumulating cells (p= 0.0013, Ctrl n=16 gonads; *org1* OE n=18 gonads, Mann-Whitney Test). Scale bars, 10 um. n≥3 trials.

The accumulation of Islet in response to *org1* parallels the transcriptional cascade that specifies embryonic alary muscle specification (Boukhatmi et al., 2014). We thus hypothesized that *islet* may also play a role in continued specification of niche identity in addition to the role we previously found during assembly (Anllo & DiNardo, 2022). To test this hypothesis, we quantified the number of Fas3 accumulating niche cells in *islet* mutants compared to controls. Indeed, we counted fewer Fas3 accumulating cells in *islet* mutants, supporting our hypothesis (Figure 6A-6C). This reduction in Fas3 accumulating cells is less striking than the complete loss of Fas3 accumulation in the majority of *org1* mutant gonads (Figure 2A-B; 3E), suggesting a more prominent role for *org1* in continued niche specification.

**Figure 6.**
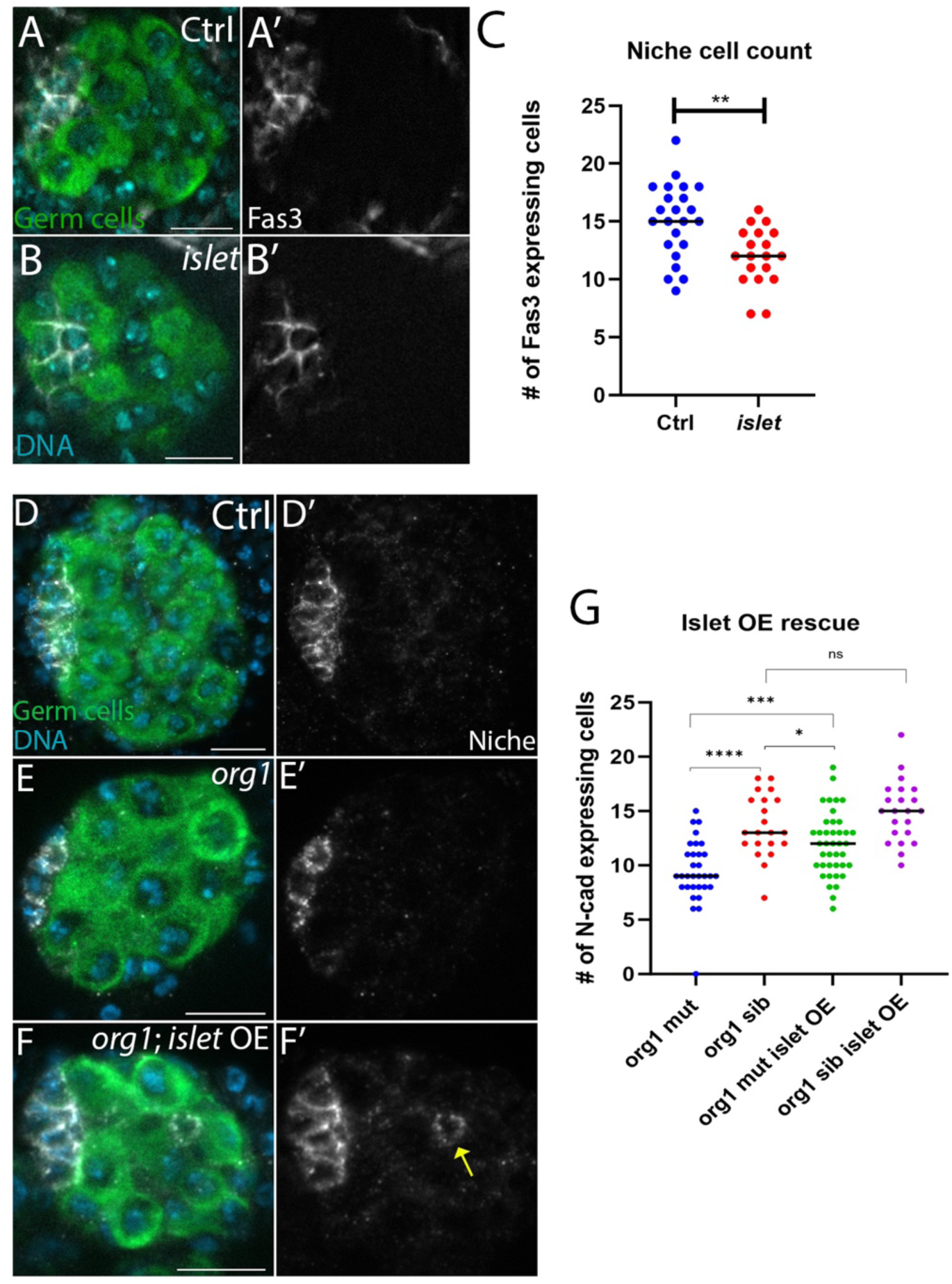
*islet* regulates niche cell specification downstream of *org1*. Germ cells (Vasa, green), DNA (Hoechst, blue), Niche cells (white). Scale bars, 10 um. (A, A’) Control gonads with niche (Fas3, white) at the anterior. (B, B’) *islet* mutant gonad with disrupted niche (Fas3, white). (C) Quantification of niche cells per gonad in *islet* mutants (p= 0.0019, Ctrl n=23 gonads; *islet* n=19 gonads, Mann-Whitney Test). (D, D’) Control gonads with N-cad (white) and Islet (red) accumulating in niche cells. (E, E’) *org1* mutant gonad with fewer niche cells accumulating N-cad at the anterior. (F, F’) An *org1* mutant with *islet* overexpression in somatic gonadal cells had more niche cells than *org1* mutants, and a displaced niche cell (yellow arrow). (G) Number of niche cells per gonad for *org1* rescue with *islet* overexpression experiments (*org1* mut vs *org1* sib, p<0.0001, *org1* mut n=32 gonads; *org1* sib n=21 gonads. *org1* mut vs *org1* mut *islet* OE, p=0.0005, *org1* mut n=32 gonads; *org1* mut *islet* OE n=40 gonads. *org1* sib vs *org1* mut *islet* OE, p=0.0338, *org1* sib n=21 gonads; *org1* mut *islet* OE n=40 gonads. *org1* sib vs *org1* sib *islet* OE, not significant, *org1* sib n=21 gonads; *org1* sib *islet* OE n=22 gonads) Mann-Whitney Tests. Scale bars, 10 um. n≥3 trials.

To further support a role for *islet* in niche specification downstream of *org1,* we asked whether we could rescue *org1* mutant niche cell number with *islet* overexpression. We overexpressed *islet* with our SGP lineage-specific driver *six4-*gal4. In gonads with both the *org1* mutation and SGP-specific restoration of *islet*, we saw increased numbers of niche cells compared to *org1* mutants alone (Figure 6G-6J’), supporting an ability to rescue niche cell identity. While niche cell number increased with *islet* overexpression, this number was still not fully rescued to control levels, and *islet* overexpression in control genetic backgrounds does not significantly increase niche cell number (Figure 6G). These results suggest that while *islet* clearly contributes to niche identity, its expression alone is not fully sufficient to induce niche cell fate. The partial rescue also suggests that *islet* is not the only relevant *org1* target for continued specification. Taken together, our results support a role for *islet* in niche cell specification and suggest that it acts downstream of *org1* in this role.

## Discussion

An impressive body of work has explored stem cell and niche functions (Anllo & DiNardo, 2022; Kaestner et. al., 2019; Losick et al., 2011; Morrison & Spradling, 2008; Sato et al., 2011; Vida et al., 2024; Warder et. al., 2024; Wei et al., 2023; Yamashita et al., 2005), yet how niche cells acquire their identity and are positioned to properly regulate resident stem cells is largely unknown. Prior to our work, Notch signaling and a downstream transcriptional regulator, *traffic jam,* were identified as inputs to *Drosophila* testis niche identity (Kitadate & Kobayashi, 2010; Okegbe & DiNardo, 2011; Wingert & DiNardo, 2015). These inputs are first induced by early contact with the developing gut (Okegbe & DiNardo, 2011). Our previous work studying extrinsic signals important for niche establishment has linked niche assembly with niche function in this system (Anllo & DiNardo, 2022), and prior work has linked niche cell identity and assembly (Wingert & DiNardo, 2015). Here, we have identified a novel regulator that further supports a connection between the processes of niche identity and assembly, the Tbx1 ortholog *org1*. We show that *org1* is expressed in response to Slit and FGF signals, which we previously identified as niche assembly cues from neighboring visceral muscle (Anllo & DiNardo, 2022; this work). In addition, we have defined *islet* as a regulator of niche identity operating downstream of *org1* (Fig 7, model). The combination of these factors suggests that niche assembly and identity are controlled through overlapping processes. Our work addresses a knowledge gap in mechanisms establishing a functional stem cell niche.

**Figure 7.**
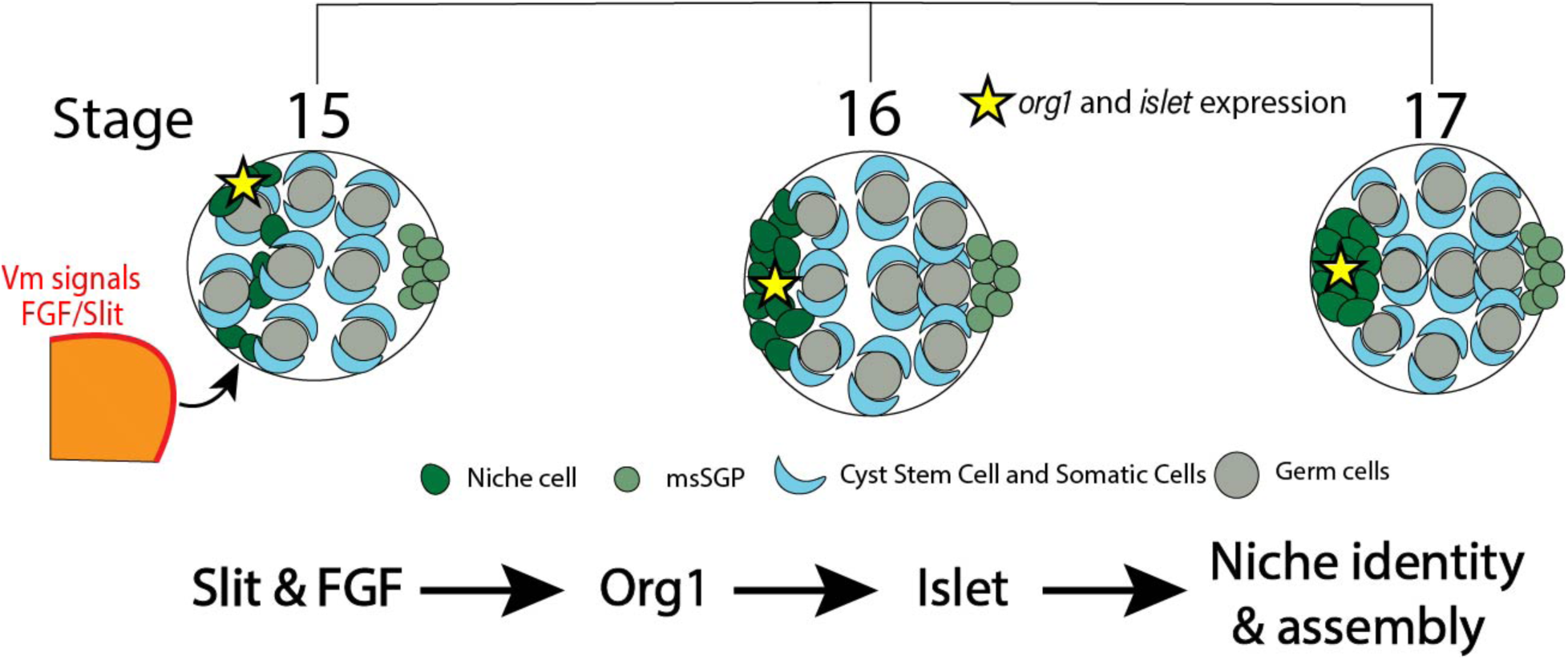
Predicted model of establishment of niche identity and assembly. Diffusible signals Slit and FGF regulate *org1*. The Visceral muscle (Vm) is a neighboring tissue source of both signals. *org1* functions to regulate *islet* for niche identity and assembly.

### Testis niche identity and assembly are linked

Our previous work showed that the Visceral muscle signals Slit and FGF are required for niche assembly. When niche assembly was compromised, niche function was also disrupted even though these manipulations did not directly affect niche identity (Anllo & DiNardo, 2022). Previous work suggested a connection between niche identity and assembly (Wingert & DiNardo, 2015). In response to the niche specification signal Notch, the Maf transcription factor *traffic jam* (*tj*) is downregulated. *traffic jam* mutants specify additional niche cells that do not express all niche identity markers, and as a result do not assemble at the anterior (Wingert & DiNardo, 2015). Interestingly, activating other requirements of niche identity such as Bowl places the additional *tj* mutant niche cells at the gonad anterior (Wingert & DiNardo, 2015; DiNardo et al., 2011). This suggests that reduction of *traffic jam* alone is not sufficient for niche assembly at the anterior, supporting a role for additional regulators specifying niche cells capable of assembly. Perhaps the signals necessary for anterior assembly (Slit and FGF) are unable to be received by ectopic niche cells in *tj* mutants.

We have shown that niche assembly signals Slit and FGF are important for regulating the novel niche identity regulator *org1*. Gonads with compromised *org1* show disruptions to markers of niche identity, including complete loss of Fas3, reduction of *hh*, loss of *upd1* expression, and fewer cells accumulating N-cadherin. The few niche cells that do accumulate N-cad exhibit assembly defects in which a portion of these cells do not arrive at the anterior (this work). The loss of continued niche specification after earlier Notch signaling might account for incomplete assembly when *org1* is disrupted. *traffic jam* mutants show additional cells expressing some niche identity markers, and a portion of these cells are also located in positions away from the gonad anterior (Wingert & DiNardo, 2015). Together with work presented here, these data link proper specification of niche identity with anterior assembly. While these data support that niche identity is important for aspects of niche assembly, some niche cells still assemble anteriorly in these instances when continued specification is disrupted. Perhaps differential adhesive sorting enables partially specified pro-niche cells at the gonad anterior to assemble.

Interestingly, the loss of *slit* and *fgf* regulators do not completely mimic phenotypes seen in *org1* mutants. For instance, *slit* and *fgf* signaling mutants are capable of accumulating Fas3, whereas *org1* mutants do not. We detect residual Org1 accumulation in niche cells from *slit* and *fgf* mutants (Figure 1) suggesting that additional regulators are impacting *org1* expression. We suspect that while Slit and FGF both regulate *org1*, Slit and FGF signaling have a primary role in facilitating niche cell anterior placement, while *org1* has a primary function in continued specification of niche cell identity.

### *islet* has dual roles in niche identity and assembly downstream of *org1*

To further link niche identity and assembly, we present an exciting new role for the transcription factor *islet* in niche specification in addition to the role we previously revealed for *islet* in niche assembly. Together with our previous study, this work reveals that Slit and FGF signals induce both *org1* and *islet* expression in niche cells (Anllo & DiNardo, 2022; this work). The visceral muscle (Vm) is an exciting candidate source of the Slit and FGF ligands inducing *org1*, as previous work revealed Vm as an important tissue supplying these signals for niche assembly (Anllo & DiNardo, 2022). In addition, we show that *org1* regulates *islet* in the niche, as *org1* mutant niches lack Islet, and niche cell numbers are increased by overexpressing *islet* in SGPs in an *org1* mutant background. These data support the model of a transcriptional cascade regulating *islet* first through Slit and FGF signals inducing *org1,* which in turn induces *islet* expression (Figure 7). *org1* has a more prominent role in continued niche specification than *islet*, as *org1* mutants often lack Fas3 accumulation (Figure 2A-B; 3E), whereas *islet* mutants have fewer Fas3 positive niche cells (Figure 6B). This result, along with the finding that *islet* overexpression only partially rescues niche cell number (Figure 6E), suggests that other genes act downstream of *org1* in addition to *islet*. Interesting questions that remain include identifying these additional *org1* targets that contribute to continued specification, and revealing how *islet* influences the number of niche cells that accumulate Fas3 and N-cad.

The detection of dispersed, unassembled niche cells in our *six4*VP16-gal4 driven *org1* RNAi was unexpected. It is reasonable to suggest earlier and stronger knockdown driving *org1* RNAi with the enhanced *six4*VP16-Gal4 driver compared to that observed with *six4*-Gal4. Since we only observed assembly defects under the presumed stronger conditions, we infer that *org1* has a primary role in niche cell identity and a secondary role in niche assembly.

### *org1* regulates testis niche identity

In alignment with Tbx1 orthologs and their conserved role in specifying mesodermal cell types, we found it fitting to test if *org1* had a role in specifying the mesodermally-derived niche cells. Using the niche identity markers Fasciclin3 (Fas3), *unpaired1* (*upd1),* and *hedgehog* (*hh*), we saw that *org1* mutants were unable to properly accumulate these markers (Le Bras & Van Doren, 2006; Okegbe & DiNardo, 2011). *hh* and its expression have been linked to niche assembly downstream of Notch (Wingert & DiNardo, 2015). Using the *hh*-lacZ reporter, we detected high *hh* expression in the niche, and low expression in SGPs outside the niche (Figure S2). While we find that *hh* expression is not solely restricted to the niche in the embryonic gonad as it is in the adult (Amoyel et. al., 2013), our ability to detect higher *hh* expression in niche cells enables us to use its upregulated expression as a reliable marker for niche identity.

There are residual niche cells in *org1* mutants, which suggests that other inputs of niche identity are partly active. It has been previously described that Notch signaling is necessary for specifying niche cells (Kitadate & Kobayashi, 2010; Okegbe & DiNardo, 2011). The average number of niche cells in Notch mutants is far fewer than we see in *org1* mutants, which suggested to us that *org1* is either one of multiple effectors downstream of Notch or involved in a parallel niche specification mechanism. One of the effects of Notch signaling in specifying niche cells is to downregulate Traffic jam (Tj) accumulation in these cells. Previously, it was not known when downregulation of *traffic jam* began in prospective niche cells (Wingert & DiNardo, 2015). We detected downregulation of *tj* coinciding with the beginning of *org1* expression in prospective niche cells after gonad coalescence. Since *org1*-expressing cells had lowered levels of Tj, we considered whether *org1* might repress Tj. However, we have shown that Tj is not regulated by *org1* (Figure S2). Alternatively, Tj might block *org1* expression. This relationship could explain the higher accumulation or Org1::GFP we detect in SGPs with reduced Tj (Figure 1B-B’’, yellow arrows). Future work should assess whether Tj regulates *org1* downstream of Notch.

Our work has established *org1* as a novel regulator of identity and assembly during testis niche establishment. Multiple factors identified in the *Drosophila* testis stem cell niche have translated into other systems, including a role for Notch in niche specification (Zamfirescu et al., 2022). Because of the numerous cell biological concepts elucidated in this niche that have translated to other systems, our work lays the foundation for future studies to investigate a conserved capacity of Tbx1 to specify gonadal mesoderm or niche cell fate.

### Study limitations

While we reveal a role for *org1* in continued niche specification during development, our genetic manipulations do not survive past embryogenesis. This precludes our ability to assess later effects of losing *org1* during development. This work importantly reveals a role for *org1* during initial niche formation, and future work will be required to study a role for *org1* in adult niche maintenance. Finally, while assessing double mutant animals would be one way to support evidence of *islet* downstream of *org1*, genetic odds of conducting this experiment are not feasible.

## Methods

### Embryonic gonad dissection and immunostaining

Embryos were collected for 2 hours and then aged for 20-22 hours in a humidified chamber at 25 C to stage 17. Embryos were then sorted for mutant and control genotypes using fluorescently marked balancer chromosomes (FM7DfdYFP or TM6DfdGFP). Sorted embryos were dissected and immunostained as previously described (Anllo et. al. 2019). Dissected tissue was incubated at 4C overnight with primary antibodies. Secondary antibodies were used at 3.75 ug/mL (Alexa488, Cy3, or Alexa647; Molecular Probes; Jackson Immunoresearch) for 1.5-2 hours at room temperature. DNA was stained with Hoechst33342 (Sigma) at 0.2-0.3 ug/mL for 6 minutes.

We used goat anti Vasa 1:800 (Santa Cruz Biotech -Discont.); rabbit anti Vasa 1:5000 (BosterBio DZ41154); mouse anti Fasciclin III 1:50 (DSHB AB_528238); rat anti N-cadherin 1:20 (DSHB AB_528121); rat anti E-cad 1:20 (DSHB AB_528120); rabbit anti Stat92E 1:200 (Flaherty et al., 2010); rabbit anti RFP 1:500 (Abcam ab62341); mouse anti Islet 1:200 (DSHB AB_528313); guinea pig anti Traffic jam 1:10,000 (Gift from D. Godt); rat anti Org-1 1:200 (gift from C. Schaub); mouse anti Gamma Tubulin 1:200 (Sigma AB_477584); chicken anti Beta-gal 1:1000 (Abcam ab9361-250); mouse anti Patched 1:50 (DSHB AB_528441); and chicken anti GFP 1:1000 (Aves labs AB_2307313).

Tissue was imaging using a Zeiss AxioImager.Z1 microscope equipped with and ApoTome.2, X-cite Xylis fluorescent light source, and Axiocam 705 mono camera, using 40x 1.2 N.A. plan apo lens. Org1::GFP reporter images in Stage 15 embryos were obtained with a Crest X-Light V3 spinning disk confocal on an Olympus IX83 platform with a 60x 1.3 N.A. plan apo silicone oil immersion objective.

### Visualization of Stage 15 embryonic gonads

Embryos were collected on grape juice agar plates supplemented with a dab of fresh yeast paste. After allowing adult flies to lay fertilized eggs for 16 h, the embryos were transferred to a nylon mesh strainer by rinsing with de-ionized water and using a paintbrush. Embryos were de-chorionated with 50% bleach for no longer than 5 min. Embryos were fixed in a glass vial at 23 degrees C in a 1:1 mixture of 4% paraformadelhyde (PFA) in PBS:heptane for 18 min while on a nutator (Mitchison and Sedat, 1983). The PFA was removed and replaced with an equal volume of 100% methanol. The vial was vigorously shaken for 30-60 seconds to devitellinize the embryos. Heptane was then removed, and devitellinized embryos were washed with 100% methanol three times. Embryos were washed once with 50% methanol in phosphate buffered saline (PBS), then again with 100% PBS with 0.1% Triton for gentle rehydration. Embryos were blocked for at least 1 hour in 4% Normal Donkey Serum, prior to immunostaining and imaging as described above. Stage 15 embryos selected for imaging based on Campos-Ortega and Hartenstein (1985).

#### Drosophila strains

All *Drosophila* lines used are listed in the Key Resources Table. *org1[OJ487]* is a null allele lacking the translation initiation codon and T-box domain (Schaub et al., 2012). Org1::GFP is a bacterial artificial chromosome (BAC) including the *org1* coding sequence and nearby genetic regulatory elements upstream of an in-frame GFP sequence (Kudron et al., 2018). We have confirmed that this BAC rescues *org1* lethality (data not shown). *slit[2]* is a null allele with no detectable protein product (Battye et. al., 2001; Nusslein-Volhard et. al., 1984). To remove *pyr* and *ths* FGF ligands together, we used a small chromosomal deficiency, Df(2R)BSC25, which completely deletes the genes encoding both ligands that bind the Heartless FGF receptor (Stathopoulos et al., 2004). When generating *org1[OJ487]*, *upd1*-Gal4 recombinant chromosomes, we screened for homozygous lethality and the ability to drive *upd1*-Gal4, UAS-redStinger expression in larval tissues.

### Identification of male gonads, and the niche

We identified male gonads by the presence of male specific SGPs at the posterior (De Falco et. al., 2003) which were visualized with Vasa (BosterBio) or six4moeGFP (Sano et al., 2012). Additionally, male gonads can be identified by the fact that they have more proliferative germ cells and therefore have more germ cells than female gonads (Anllo et. al., 2019; Wawersik et. al., 2005). Niches in stage 17 embryos were identified via immunofluorescence by niche cell specific accumulation of Fasciclin 3 (DSHB AB_528238), or enrichment of N-cadherin (DSHB AB_528121) or E-cadherin (DSHB AB_528120).

### Characterization of niche phenotypes

When assessing morphology and proper assembly of the niche, we characterized a niche as either “assembled” or “unassembled”. “Assembled” niches show all niche cells contacting one another at the gonad anterior and have clearly smoothened cell boundaries around the niche periphery. “Unassembled” niches have niche cells anterior and at the gonad periphery, but with irregular boundaries, and/or have dispersed niche cells that are not contacting the anterior cluster of niche cells.

### Rigor and Reproducibility

All experiments were conducted at least in triplicate (n≥3 trials) with sufficiently large sample sizes that previously enabled detection of significant differences (significance level of less than or equal to 0.05%). Genetic controls were always used, comparing perturbed conditions to the most genetically similar control background feasible.

### Quantification of niche cell count

When quantifying the number of cells that express N-cad, Fas3, or any other niche marker, we use the ImageJ Cell Counter Plugin to record cells that we counted manually. A niche cell will accumulate Fas3 when in contact with other niche cells and will accumulate N-cad on all interfaces. Niche cells that are counted with either Fas3 or N-cad are co-labeled with a nuclear marker such as Traffic jam or Hoechst, to enable clear identification of distinct cells. Dispersed niche cells that accumulate Fas3 do not show polarized enrichment to specific cell interfaces.

### Quantification of normalized niche cell expression/accumulation

When quantifying accumulation of proteins and reporters of gene expression, we used ImageJ software to measure mean gray value fluorescence intensity within regions of interest (ROIs). We selected ROIs including a circular region within somatic cell boundaries, using either Fas3 or N-cad immunofluorescence to delineate niche cell boundaries and Hoechst dye to delineate nuclei. ROIs were in a single Z plane in which the relevant cell was in focus. For all experiments except *upd* measurements, 3 niche cell ROIs were measured from each gonad. For *upd* measurements, 5 niche cell ROIs were measured per gonad. To quantify background fluorescence, an ROI was selected to encompass the unlabeled region of a single germ cell within each gonad, and a separate background ROI drawn from a region where no tissue was present. Background-subtracted fluorescence values for each niche cell were normalized, dividing by the background-subtracted value from the neighboring unlabeled germ cell for the respective gonad. Relative fluorescence values for each niche cell were plotted. Mann-Whitney tests were used to evaluate comparisons.

### Quantification of fluorescence at cell junctions

To quantify fluorescence intensity of F-actin between cell interfaces, we used ImageJ software to measure mean gray value fluorescence intensity within regions of interested (ROIs). ROIs were either niche-niche interfaces or niche-GSC interfaces. We selected 4-pixel-wide segmented line ROIs between niche cells and between niche cells and GSCs. 2-4 niche-niche and niche-GSC interfaces were measured in every gonad using N-cad to visualize each interface. We measured the mean fluorescence intensity of our Ftractin-tdTomato blindly with N-cad as a boundary. To estimate background fluorescence, a separate ROI was drawn from a region where no tissue was present. Background-subtracted fluorescence values for each niche cell were normalized, divided by the background-subtracted average value of all cell interfaces. Relative fluorescence values for each interface were plotted. Mann-Whitney tests were used to evaluate comparisons.

### Quantification of Stat accumulation

To quantify Stat accumulation, we stained gonads with a Stat antibody (E. Bach, 1:200) and used ImageJ to measure the mean gray value fluorescence intensity within regions of interest (ROIs). During dissections and staining, we used the phosphatase inhibitor PhosStop (Sigma – Cat#4906845001). We selected ROIs including a circular region to sample germ cells, using Vasa immunofluorescence as a marker to delineate cell boundaries. Whole germ cells were measured in a single Z plane in which the relevant cell was in focus. We selected ROIs by only using the germ cell marker to mark sample regions to avoid biased observations towards brighter or dimmer fluorescence of Stat. For each gonad, we sampled all GSCs we could clearly identify and 3-7 neighboring germ cells in the second tier, further from the germline stem cells. After background subtraction, we measured the ratio of Stat accumulation within each GSC relative to the neighboring germ cell average for that gonad. Relative Stat enrichment values were plotted for each GSC. We obtained measurements on mutants for *org1* and their sibling controls, and *org1* RNAi knockdowns with their sibling controls. Mann-Whitney tests were used to evaluate comparisons.

### Quantification of STAT+ GSCs

To determine whether a germ cell adherent to the niche was STAT+, we first counted the number of germline stem cells (GSCs) directly adjacent to N-cadherin labeled niche cells. We then quantified the mean fluorescence intensity from each of these adherent germ cells as described above, measuring mean STAT enrichment for each adherent germ cell in *org1* RNAi knockdowns and sibling control animals. We established a threshold STAT+ fluorescent value, measured as one standard deviation below the relative mean fluorescence intensity of controls. For controls, the relative mean fluorescence intensity was 2.76, with a standard deviation of 0.97, therefore any adherent germ cells with a mean fluorescence intensity above 1.79 were counted as STAT+ in both control and *org1* RNAi knockdown gonads. Data were imported into Prism v10.0 to generate scatterplots where each dot represents a gonad. We plotted the number of STAT+ GSCs per gonad, and separately the percentage of STAT+ GSCs per gonad for *org1* RNAi and sibling control conditions. A Mann–Whitney test was used to evaluate the comparison.

### Centrosome anchoring quantification

Centrosome position was visualized with immunofluorescence against Gamma tubulin to label pericentriolar material. GSCs were scored for centrosome position if they had already undergone centrosome duplication, indicating that they had advanced at least to the G2 stage of the cell cycle (Chen et. al., 2018). We quantified how often one of the two centrosomes was positioned closer to the adjacent niche, marked by N-cad, than to other neighboring cells. GSCs with a centrosome located near the niche-GSC interface were scored as appropriately aligned. GSCs with both centrosomes positioned away from the niche were scored as misaligned. Significance was assessed with Fisher’s exact test.

## Key Resources Table

**Key Resources Table:**
| Reagent Type | Designation | Source | Identifiers |
| --- | --- | --- | --- |
| Genetic reagent (D. melanogaster) | org1[OJ487]/FM7DfdYFP or Y | Bloomington Drosophila Stock Center | Schaub et. al., 2012; RRID: BDSC_91555 |
| Genetic reagent (D. melanogaster) | org1 RNAi (II) | Vienna Drosophila Stock Center | v104393 |
| Genetic reagent (D. melanogaster) | UAS-org1 | Bloomington Drosophila Stock Center | Schaub et. al., 2012; RRID: BDSC_92971 |
| Genetic reagent (D. melanogaster) | six4-gal4/CyoDfdYFP; Dr/Tm3,sb | Anllo et al., 2019 |  |
| Genetic reagent (D. melanogaster) | six4VP16-gal4/CyoDfdYFP; Dr/TM3,Sb | Warder et al., 2024 |  |
| Genetic reagent (D. melanogaster) | w[*] ; P{w+ UAS-islet} on III | Bloomington Drosophila Stock Center | RRID:BDSC_39662 |
| Genetic reagent (D. melanogaster) | ry[506], P{ry+}hhLacZ | Bloomington Drosophila Stock Center | RRID:BDSC_5530 |
| Genetic reagent (D. melanogaster) | P-Dsix4-eGFP::Moesin | R. Lehmann | Sano et. al., 2012 |
| Genetic reagent (D. melanogaster) | Pbac{org-1-GFP.FPTB}VK00033/TM6C, Sb[1] | Bloomington Drosophila Stock Center | Kudron et al., 2018 |
| Genetic reagent (D. melanogaster) | six4>ftractin tdTomato on II | Stephen DiNardo | RRID: BDSC_58989<br>UAS-ftractin tdTomato |
| Genetic reagent (D. melanogaster) | Df(2R)BSC25 – *fgf* mutant | Bloomington Drosophila Stock Center | RRID: BDSC_6865 |
| Genetic reagent (D. melanogaster) | *slit[2]* | Bloomington Drosophila Stock Center | RRID: BDSC_3266<br>(Battye et. al., 2001; Nusslein-Volhard et. al., 1984) |
| Genetic reagent (D. melanogaster) | tupex4 | S. Campuzano | FBal0216723;<br>(Stathopoulos et al., 2004). |
| Genetic reagent (D. melanogaster) | *nos*-lifeactin-tdTomato | B. Lin; R. Lehmann | (Lin et al., 2020) |
| Genetic reagent (D. melanogaster) | *upd1*-Gal4 | Bloomington Drosophila Stock Center | RRID: BDSC_26796 |
| Genetic reagent (D. melanogaster) | UAS-redStinger | Bloomington Drosophila Stock Center | RRID: BDSC_8547 |
| Antibody | Goat polyclonal anti-Vasa | Santa Cruz Biotechnology | RRID:AB_793880<br>Discontinued |
| Antibody | Mouse monoclonal anti Fasciclin III | Developmental Studies Hybridoma Bank | RRID:AB_528238 |
| Antibody | Rabbit polyclonal anti STAT92E | E. Bach |  |
| Antibody | Rabbit polyclonal anti RFP | Abcam | RRID:AB_945213 |
| Antibody | Mouse monoclonal anti Islet | Developmental Studies Hybridoma Bank | RRID:AB_528313 |
| Antibody | Guinea pig polyclonal anti Traffic jam | D. Godt |  |
| Antibody | Rat anti Org-1 | C. Shaub |  |
| Antibody | Mouse monoclonal anti Gamma-tubulin | Sigma | RRID:AB_477584 |
| Antibody | Rabbit Polyclonal anti Vasa | Boster Bio | DZ41154 |
| Antibody | Chicken polyclonal anti Beta-gal | Abcam | ab9361-250 |
| Antibody | Mouse Monoclonal anti Patched | Developmental Studies Hybridoma Bank | RRID: AB_528441 |
| Antibody | Rat monoclonal anti CadN | Developmental Studies Hybridoma Bank | RRID: AB_528121 |
| Antibody | Rat monoclonal anti CadE | Developmental Studies Hybridoma Bank | RRID: AB_528120 |
| Antibody | Chick polyclonal anti GFP | Aves Labs | RRID: AB_2307313 |
| Antibody | Alexafluor Secondary Antibodies (488, 647) | Molecular Probes |  |
| Antibody | Cy3 Affinipure Secondary Antibodies | Jackson ImmunoResearch | RRID: AB_2307443 |
| Chemical Compound, Drug | Hoechst | Sigma | Cat#33342; CAS Number: 875756-97-1 |
| Chemical Compound, Drug | Normal Donkey Serum (NDS) | Jackson ImmunoResearch Laboratories | RRID: AB_2337258 |
| Chemical Compound, Drug | Ringers |  | doi:10.1101/pdb.rec12409 |
| Chemical Compound, Drug | Heptane |  |  |
| Chemical Compound, Drug | Propyl-gallate | Sigma Aldrich | SKU: P3130; CAS Number 121-79-9 |
| Chemical Compound, Drug | Para-Formaldehyde (PFA), 16% | Electron Microscopy Sciences | Cat#15710 |
| Chemical Compound, Drug | Phosphatase inhibitor (PhosphaSTOP) | Roche/Sigma | Cat#4906845001 |
| Software, Algorithm | FIJI (Image J) | www.fiji.sc |  |
| Software, Algorithm | Image J | www.imagej.nih.gov/ij/ |  |
| Software, Algorithm | Graphpad Prism | Graphpad Software | RRID:SCR_002798 |
| Software, Algorithm | Axio-Vision Imaging Software | Zeiss | v4.8.1 |
| Software, Algorithm | Adobe Photoshop | https://www.adobe.com/uk/products/photoshop.html | RRID:SCR_014199 |
| Equipment | Zeiss Axioimager.Z1 | Zeiss |  |
| Equipment | ApoTome.2 | Zeiss |  |
| Equipment | Xcite Xylis | Zeiss |  |
| Equipment | 40x 1.2 NA objective | Zeiss |  |
| Equipment | Leica M165FC | Leica |  |
| Equipment | GFP Filter set ET470/<br>40x; ET525/50m | Leica 10447408 |  |

## Supplemental Figures

**Figure S1.**
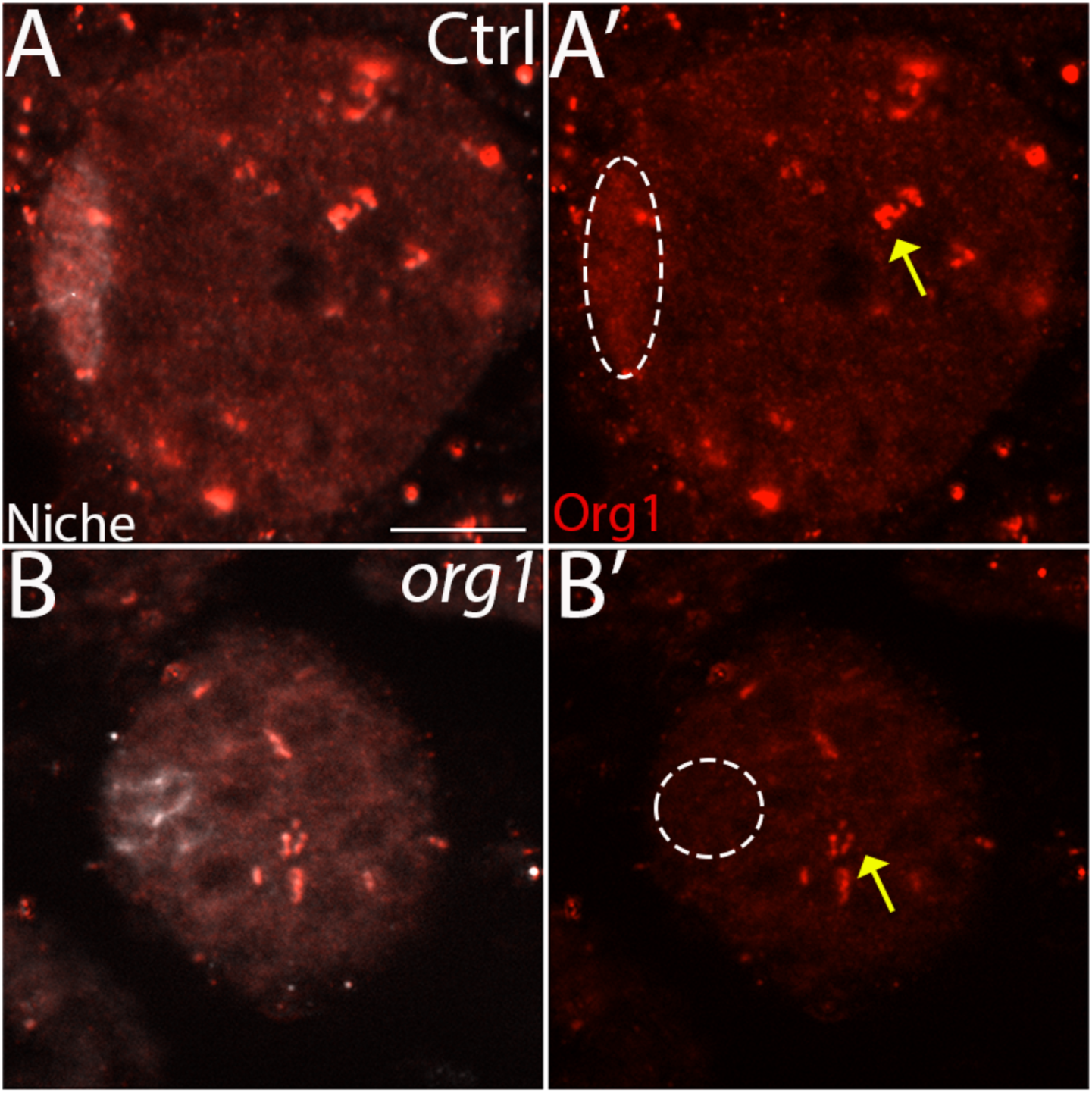
Org1 antibody validation in *org1* mutants. (A, A’) Control gonads with Org1 accumulation (red) in the niche (Fas3, white) at the anterior, along with variable punctate signal in other places in the gonad (yellow arrow). (B, B’) *org1* mutant gonads with no Org1 accumulation (red) in the niche (white) at the anterior. Note that the variable punctate signal still persists (yellow arrow) in the mutant and is thus spurious background and cannot be due to Org1. Scale bars, 10 um.

**Figure S2.**
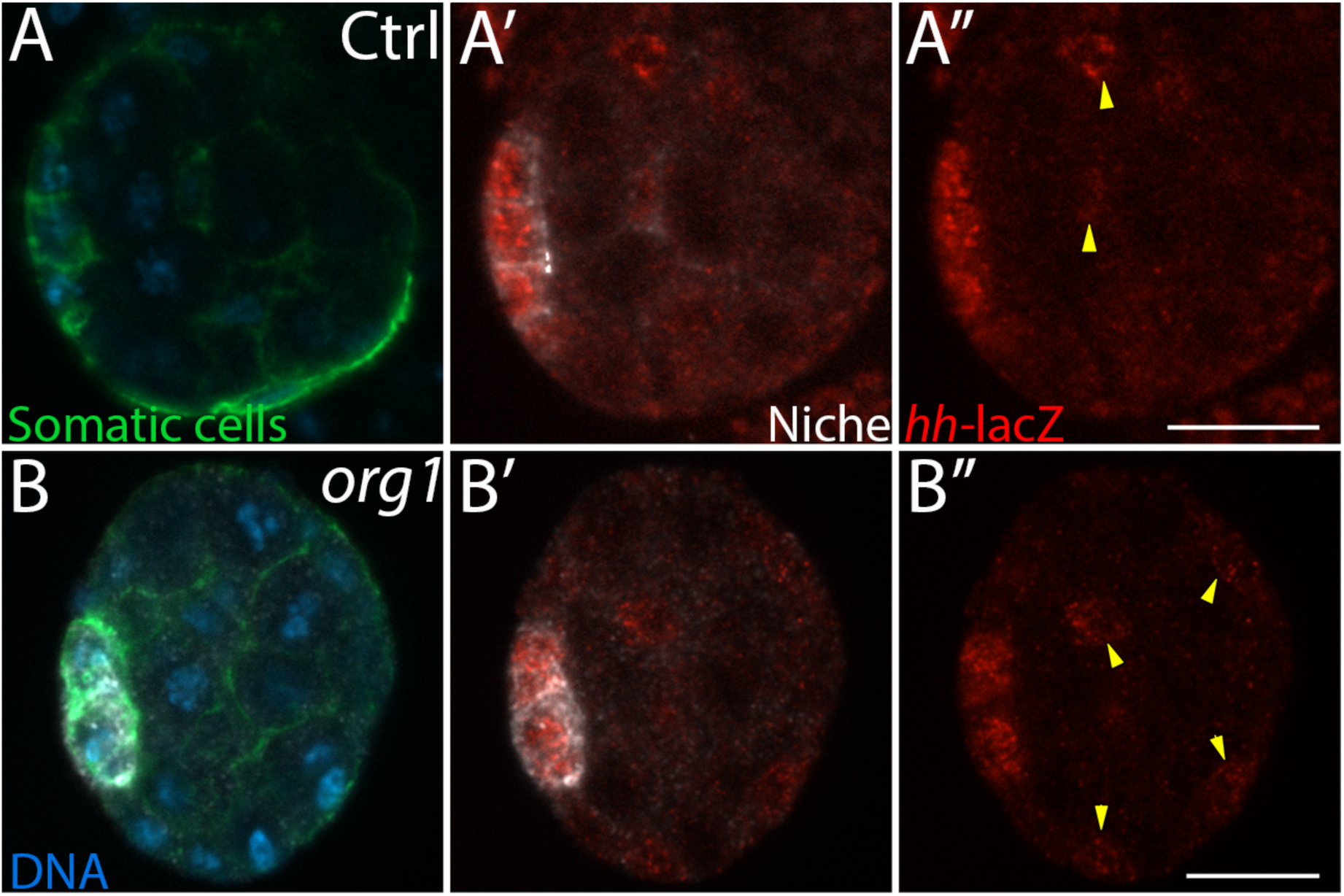
*hh* is expressed outside of the niche. Somatic cells (green), Niche (white), *hh*-lacZ (red), DNA (blue). (A-A”) *hh-*lacZ reporter can be seen at high levels in niche cells, with lower levels of expression outside of the niche (yellow arrows) in control gonads. (B-B”) *org1* mutant gonads with low *hh*-lacZ expression in the niche (quantification in Fig 2), and outside of the niche (yellow arrows).

**Figure S3.**
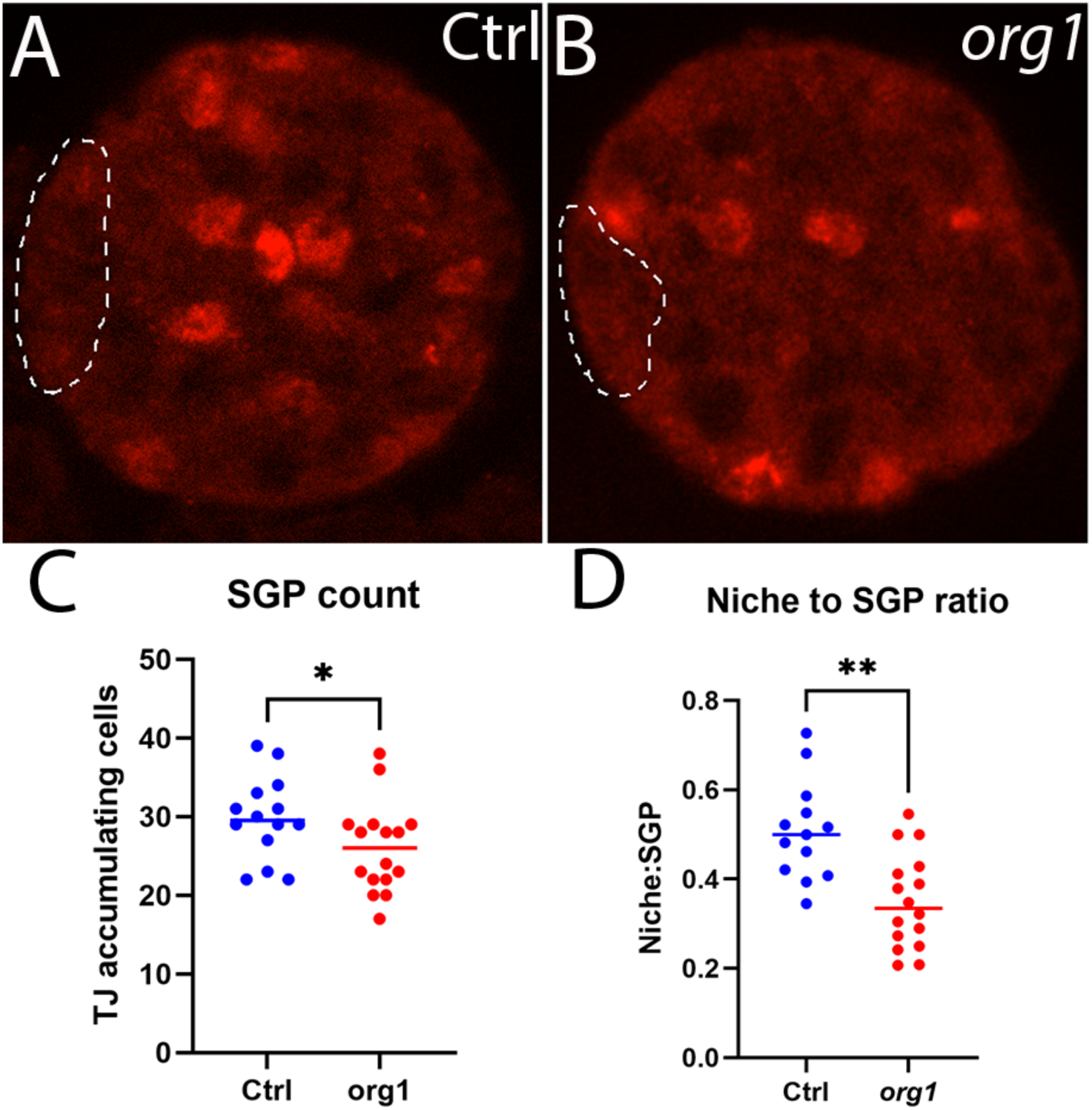
*org1* mutants exhibit a lower ratio of niche cells relative to number of SGPs. (A) Control and (B) *org1* mutant gonads with Traffic jam accumulation. Niches are outlined with dotted lines. (C) Quantification of number of Traffic jam accumulating SGPs indicated fewer SGPs in *org1* mutants versus controls (p = 0.0399, Ctrl n= 14 gonads; *org1* n=16 gonads, Mann-Whitney test). (D) Quantification of SGP to Niche cell ratio showed a lower ratio of niche cells relative to SGPs in *org1* mutants (p = 0.0010, Ctrl n=13 gonads; *org1* n=16 gonads, Mann-Whitney test). Scale bars, 10 um.

**Figure S4.**
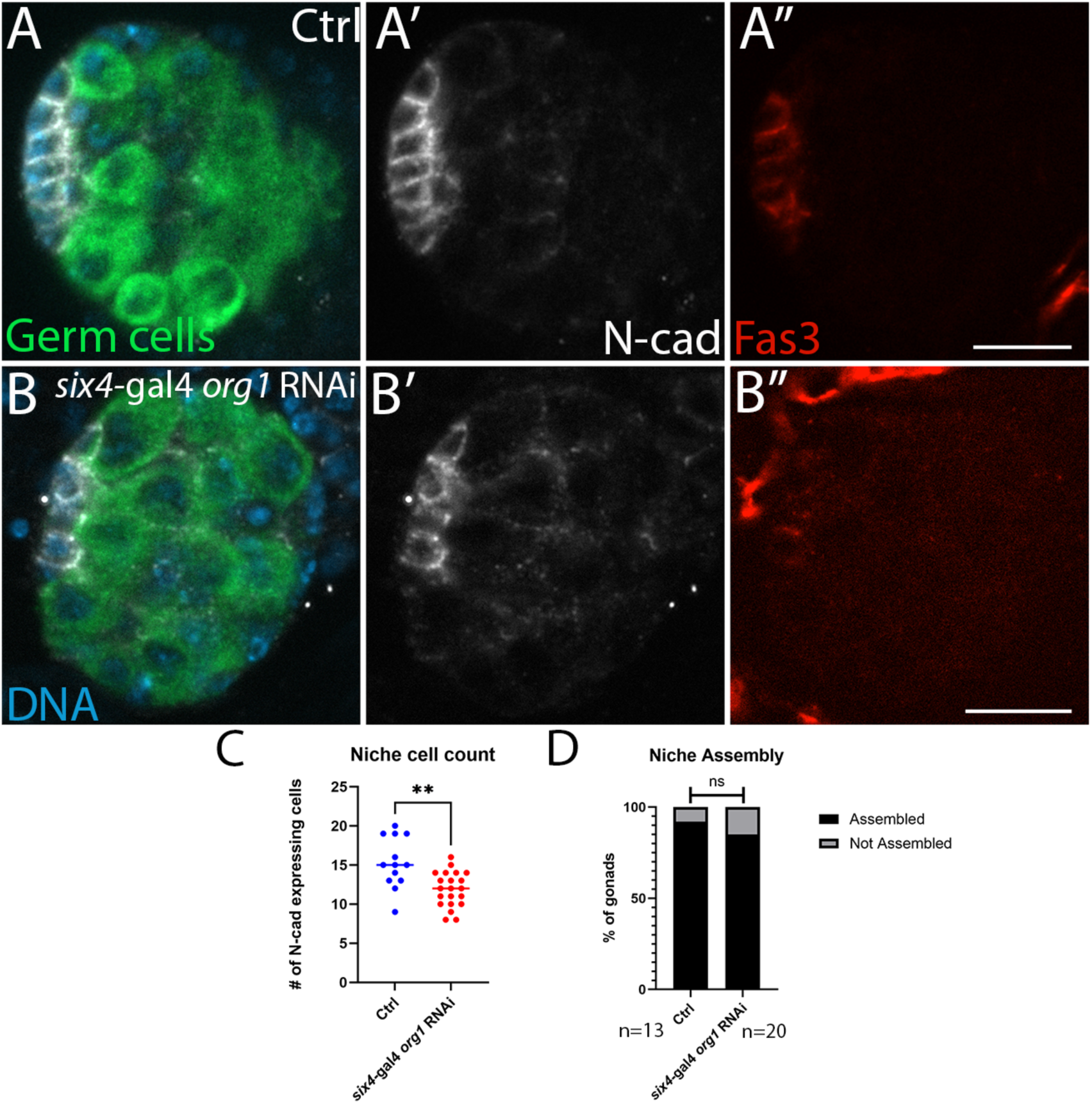
*org1* RNAi with a Gal4 driver shows identity defects but not assembly defects. Germ cells (green), N-cad (white), Fas3 (red), DNA (blue). (A-A’) Control gonads with assembled niches and clear Fas3 accumulation. (B,B’) *six4*gal4 > *org1* RNAi gonads with fewer niche cells and reduced Fas3 accumulation. (C) Quantification of niche cell count in six4-gal4 *org1* RNAi gonads (p= 0.0022, Ctrl n=13 gonads; *six4*-gal4 *org1* RNAi n=21 gonads, Mann-Whitney Test). (D) Quantification of assembly defects in six4-gal4 *org1* RNAi gonads (p> 0.99, Ctrl n=13 gonads; *six4*-gal4 *org1* RNAi n=20, Mann-Whitney Test). Scale bars, 10 um.

**Figure S5.**
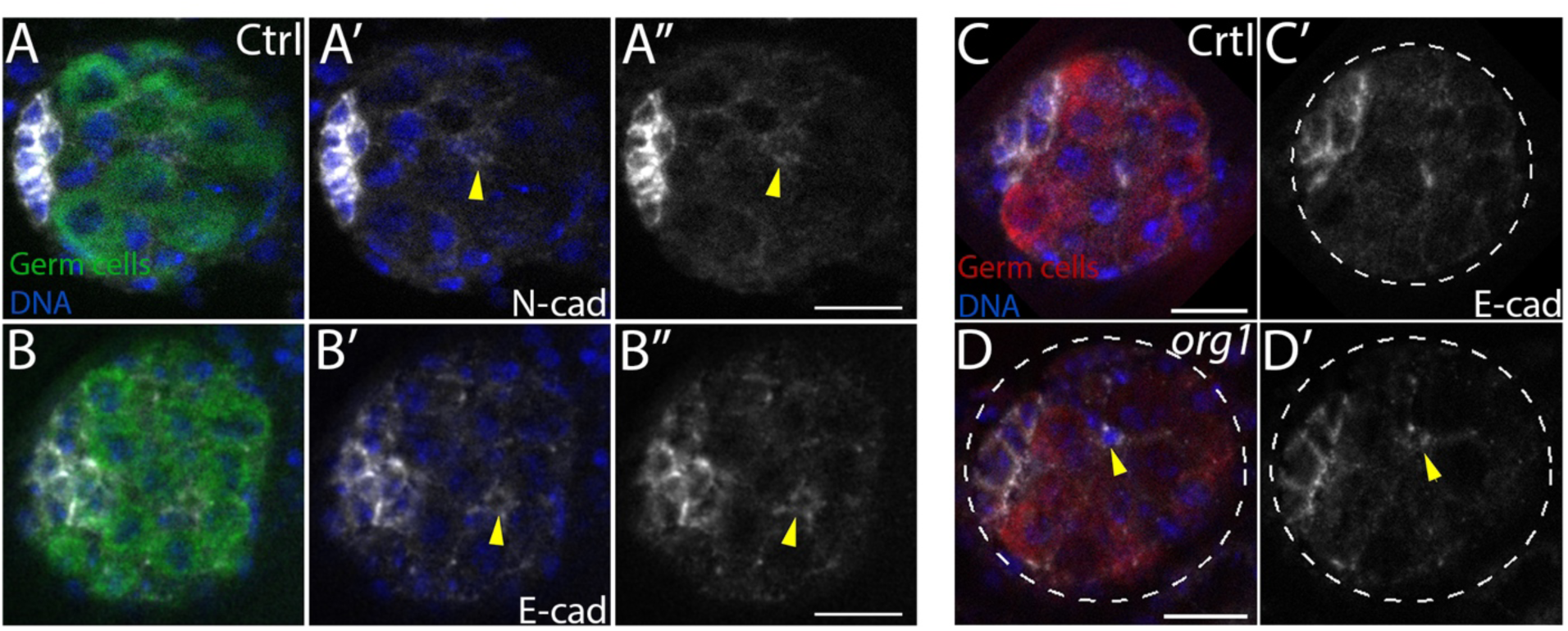
N-cad and E-cad can be found spuriously in SGPs outside of the niche. (A-B”) Control gonads immunostained to show germ cells (green) and DNA (blue). (A) N-cad (white) and (B) E-cad can be seen in an SGP (yellow arrow) outside of the niche but is not as enriched at the cell cortex as in the anterior niche. (C-D’) Germ cells (red), E-cad (white), DNA (blue). (C, C’) Control gonad with E-cad enriched in the cortex of niche cells. (D, D’) *org1* mutant gonad with E-cad enriched to similar levels in a presumptive lost niche cell outside of the anterior niche (yellow arrow). Scale bars, 10 um.

**Table S1.**
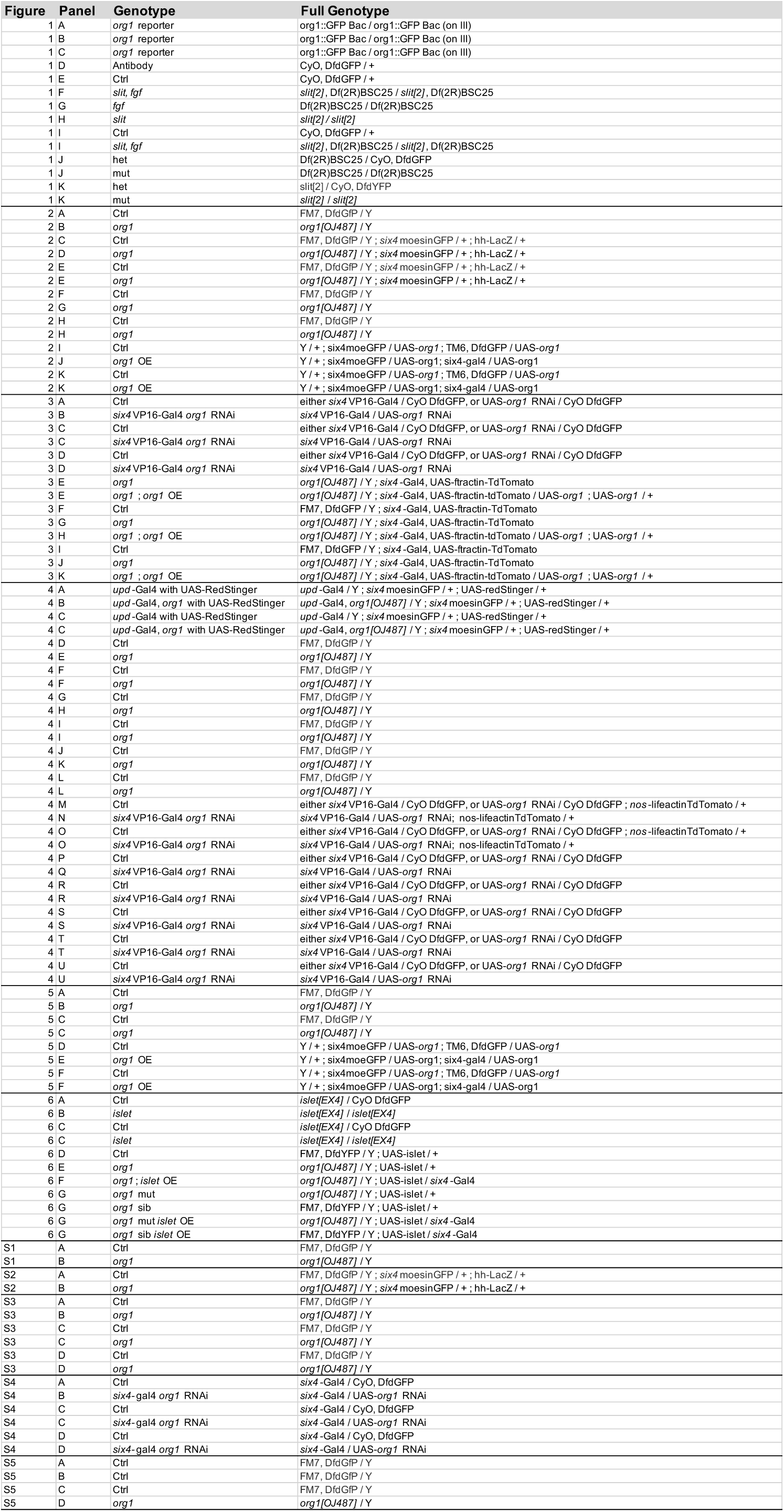
Supplementary Genotype Table.

## Acknowledgements

We acknowledge Kirklan Naumuk for technical assistance. We thank Drs. Elizabeth Ables and Beth Thompson, and the Lenhart lab for valuable feedback on the manuscript. We thank the Bloomington *Drosophila* Stock Center (NIH P40OD018537) for stocks, and D. Godt, E. Bach and C. Schaub for antibodies.

## Funding

Work was supported by NIH grants R15 GM154246 to L.A. and R35 GM136270 to S.D. Authors have no competing interests to declare.

## Author Contributions

Conceptualization, P.H. and L.A.; methodology, L.A.; validation, P.H. & L.A.; formal analysis, P.H., A.H., T.G., and L.A.; investigation P.H., T.G., and L.A.; resources S.D. and L.A.; data curation P.H. and L.A.; writing – original draft P.H.; writing – review & editing P.H., A.H., S.D., and L.A.; visualization P.H., A.H., and L.A.; supervision S.D. and L.A.; project administration S.D. and L.A.; funding acquisition S.D. and L.A.

## Notes

### Competing Interest Statement

The authors have declared no competing interest.

### Summary of Updates

This version of the manuscript has been revised to update with additional functional data in Figure 4, textual changes for clarity, and a supplementary genotype table. These changes were made in response to reviewer suggestions pre-publication. We also added an author during these revisions.

## References

1. Amoyel, M., Sanny, J., Burel, M. and Bach, E. A. (2013). Hedgehog is required for CySC self-renewal but does not contribute to the GSC niche in the Drosophila testis. Development (Cambridge) 140, 56–65.

2. Anllo, L. and DiNardo, S. (2022). Visceral mesoderm signaling regulates assembly position and function of the Drosophila testis niche. Dev Cell 57, 1009–1023.e5.

3. Anllo, L., Plasschaert, L. W., Sui, J. and DiNardo, S. (2019). Live imaging reveals hub cell assembly and compaction dynamics during morphogenesis of the Drosophila testis niche. Dev Biol 446, 102–118.

4. Battye, R., Stevens, A., Perry, R. L. and Jacobs, J. R. (2001). Repellent Signaling by Slit Requires the Leucine-Rich Repeats.

5. Boukhatmi, H., Schaub, C., Bataillé, L., Reim, I., Frendo, J. L., Frasch, M. and Vincent, A. (2014). An Org-1–Tup transcriptional cascade reveals different types of alary muscles connecting internal organs in Drosophila. Development (Cambridge) 141, 3761–3771.

6. Boyle, M. and DiNardo, S. (1995). Specification, migration and assembly of the somatic cells of the Drosophila gonad. Development.

7. Boyle, M., Bonini, N. and DiNardo, S. (1997). Expression and function of cliftin the development of somatic gonadal precursors within the Drosophila mesoderm. Development.

8. Campos-Ortega, J. A. and Hartenstein, V. (1985). The embryonic development of *Drosophila melanogaster*. Springer-Verlag, Berlin.

9. Chen, C., Cummings, R., Mordovanakis, A., Hunt, A., Mayer, M., Sept, D. and Yamashita, Y. (2018). Cytokine receptor-Eb1 interaction couples cell polarity and fate during asymmetric cell division.

10. de Cuevas, M. and Matunis, E. L. (2011). The stem cell niche: Lessons from the Drosophila testis. Development 138, 2861–2869.

11. DeFalco, T. J., Verney, G., Jenkins, A. B., McCaffery, J. M., Russell, S. and Van Doren, M. (2003). Sex-specific apoptosis regulates sexual dimorphism in the Drosophila embryonic gonad. Dev Cell 5, 205–216.

12. DiNardo, S., Okegbe, T., Wingert, L., Freilich, S. and Terry, N. (2011). Lines and bowl affect the specification of cyst stem cells and niche cells in the Drosophila testis. Development 138, 1687–1696.

13. Flaherty, M. S., Salis, P., Evans, C. J., Ekas, L. A., Marouf, A., Zavadil, J., Banerjee, U. and Bach, E. A. (2010) *chinmo* is a functional effector of the JAK/STAT pathway that regulates eye development, tumor formation, and stem cell self-renewall in *Drosophila*. Dev Cell 18, 556–568.

14. Herrera, S. C. and Bach, E. A. (2019). JAK/STAT signaling in stem cells and regeneration: From drosophila to vertebrates. Development (Cambridge) 146,.

15. Herrera, S. C., Sainz de la Maza, D., Grmai, L., Margolis, S., Plessel, R., Burel, M., O’Connor, M., Amoyel, M. and Bach, E. A. (2021). Proliferative stem cells maintain quiescence of their niche by secreting the Activin inhibitor Follistatin. Dev Cell 56, 2284–2294.e6.

16. Inaba, M., Buszczak, M. and Yamashita, Y. M. (2015). Nanotubes mediate niche-stem-cell signalling in the Drosophila testis. Nature 523, 329–332.

17. Jenkins, A. B., McCaffery, J. M. and Van Doren, M. (2003). Drosophila E-cadherin is essential for proper germ cell-soma interaction during gonad morphogenesis. Development 130, 4417–4426.

18. Kaestner, K. H. (2019). The Intestinal Stem Cell Niche: A Central Role for Foxl1-Expressing Subepithelial Telocytes. CMGH 8, 111–117.

19. Kawase, E., Wong, M. D., Ding, B. C. and Xie, T. (2004). Gbb/Bmp signaling is essential for maintaining germline stem cells and for repressing bam transcription in the Drosophila testis. Development 131, 1365–1375.

20. Kiger, A. A., Jones, D. L., Schulz, C., Rogers, M. and Fuller, M. (2001). Stem Cell Self-Renewal Specified byJAK-STAT Activation in Response to a Support Cell Cue. Science (1979) 294, 2539–2542.

21. Kitadate, Y. and Kobayashi, S. (2010). Notch and Egfr signaling act antagonistically to regulate germ-line stem cell niche formation in Drosophila male embryonic gonads. Proc Natl Acad Sci U S A 107, 14241–14246.

22. Kudron, M. M., Victorsen, A., Gevirtzman, L., Hillier, L. W., Fisher, W. W., Vafeados, D., Kirkey, M., Hammonds, A. S., Gersch, J., Ammouri, H., et al. (2018). The modern resource: genome-wide binding profiles for hundreds of Drosophila and Caenorhabditis elegans transcription factors. Genetics 208, 937–949.

23. Le Bras, S. and Van Doren, M. (2006). Development of the male germline stem cell niche in Drosophila. Dev Biol 294, 92–103.

24. Leatherman, J. L. and Di Nardo, S. (2008a). Zfh-1 controls somatic stem cell self-renewal in the Drosophila testis and nonautonomously influences germline stem cell self-renewal. Cell Stem Cell 3, 44–54.

25. Leatherman, J. L. and Di Nardo, S. (2008b). Zfh-1 controls somatic stem cell self-renewal in the Drosophila testis and nonautonomously influences germline stem cell self-renewal. Cell Stem Cell 3, 44–54.

26. Leatherman, J. L. and Dinardo, S. (2010). Germline self-renewal requires cyst stem cells and stat regulates niche adhesion in Drosophila testes. Nat Cell Biol 12, 806–811.

27. Lee, S., Zhou, L., Kim, J., Kalbfleisch, S. and Schöck, F. (2008). Lasp anchors the Drosophila male stem cell niche and mediates spermatid individualization. Mech Dev 125, 768–776.

28. Lenhart, K. F. and DiNardo, S. (2015). Somatic Cell Encystment Promotes Abscission in Germline Stem Cells following a Regulated Block in Cytokinesis. Dev Cell 34, 192–205.

29. Li, L. and Xie, T. (2005). Stem cell niche: Structure and function. Annu Rev Cell Dev Biol 21, 605–631.

30. Li, H., Janssens, J., de Waegeneer, M., Kolluru, S. S., Davie, K., Gardeux, V., Saelens, W., David, F. P. A., Brbić, M., Spanier, K., et al. (2022). Fly Cell Atlas: A single-nucleus transcriptomic atlas of the adult fruit fly. Science (1979) 375,.

31. Li, G., Li, Z., Li, L., Liu, S., Wu, P., Zhou, M., Li, C., Li, X., Luo, G. and Zhang, J. (2023). Stem Cell-Niche Engineering via Multifunctional Hydrogel Potentiates Stem Cell Therapies for Inflammatory Bone Loss. Adv Funct Mater 33,.

32. Lin, B., Luo, J., and Lehmann, R. (2020). Collectively stabilizing and orienting posterior migratory forces disperses cell clusters in vivo. Nature Communications 11, 4477.

33. Losick, V. P., Morris, L. X., Fox, D. T. and Spradling, A. (2011). Drosophila Stem Cell Niches: A Decade of Discovery Suggests a Unified View of Stem Cell Regulation. Dev Cell 21, 159–171.

34. Marcellini, S., Technau, U., Smith, J. C. and Lemaire, P. (2003). Evolution of Brachyury proteins: Identification of a novel regulatory domain conserved within Bilateria. Dev Biol 260, 352–361.

35. Michel, M., Kupinski, A. P., Raabe, I. and Bökel, C. (2012). Hh signalling is essential for somatic stem cell maintenance in the drosophila testis niche. J Cell Sci 125, 2663–2669.

36. Mitchison, T.J. and Sedat, J. (1983). Localization of antigenic determinants in whole *Drosophila* embryos. Dev Biol 99, 261–264.

37. Morrison, S. J. and Spradling, A. C. (2008). Stem Cells and Niches: Mechanisms That Promote Stem Cell Maintenance throughout Life. Cell 132, 598–611.

38. Nusslein-Volhard, C., Wieschaus, E. and Kluding, H. (1984). Roux’s Archives of Developmental Biology Mutations affecting the pattern of the larval cuticle in Drosophila melanogaster I. Zygotic loci on the second chromosome.

39. Okegbe, T. C. and di Nardo, S. (2011). The endoderm specifies the mesodermal niche for the germline in Drosophila via Delta-Notch signaling. Development 138, 1259–1267.

40. Riechmann, V., Rehorn, K.-P., Reuter, R. and Leptin, M. (1998). The genetic control of the distinction between fat body and gonadal mesoderm in Drosophila. Development.

41. Roach, T. V. and Lenhart, K. F. (2024). Mating-induced Ecdysone in the testis disrupts soma-germline contacts and stem cell cytokinesis. Development (Cambridge) 151,.

42. Sato, T., Van Es, J. H., Snippert, H. J., Stange, D. E., Vries, R. G., Van Den Born, M., Barker, N., Shroyer, N. F., Van De Wetering, M. and Clevers, H. (2011). Paneth cells constitute the niche for Lgr5 stem cells in intestinal crypts. Nature 469, 415–418.

43. Schaub, C., Nagaso, H., Jin, H. and Frasch, M. (2012). Org-1, the Drosophila ortholog of Tbx1, is a direct activator of known identity genes during muscle specification. Development 139, 1001–1012.

44. Sheng, X. R., Posenau, T., Gumulak-Smith, J. J., Matunis, E., Van Doren, M. and Wawersik, M. (2009). Jak-STAT regulation of male germline stem cell establishment during Drosophila embryogenesis. Dev Biol 334, 335–344.

45. Sinden, D., Badgett, M., Fry, J., Jones, T., Palmen, R., Sheng, X., Simmons, A., Matunis, E. and Wawersik, M. (2012). Jak-STAT regulation of cyst stem cell development in the Drosophila testis. Dev Biol 372, 5–16.

46. Stathopoulos, A., Tam, B., Ronshaugen, M., Frasch, M. and Levine, M. (2004). Pyramus and thisbe: FGF genes that pattern the mesoderm of Drosophila embryos. Genes Dev 18, 687–699.

47. Tanentzapf, G., Devenport, D., Godt, D. and Brown, N. H. (2007). Integrin-dependent anchoring of a stem-cell niche. Nat Cell Biol 9, 1413–1418.

48. Tseng, C. Y., Burel, M., Cammer, M., Harsh, S., Flaherty, M. S., Baumgartner, S. and Bach, E. A. (2022). chinmo-mutant spermatogonial stem cells cause mitotic drive by evicting non-mutant neighbors from the niche. Dev Cell 57, 80–94.e7.

49. Tulina, N. and Matunis, E. (2001). Control of Stem Cell Self-Renewal in Drosophila Spermatogenesis by JAK-STAT Signaling. Science (1979) 294, 2546–2548.

50. Vida, G. S., Botto, E. and Dinardo, S. (2024). Maintenance of niche architecture requires actomyosin and enables proper stem cell signaling and oriented division in the Drosophila testis.

51. Walker, M. R., Patel, K. K. and Stappenbeck, T. S. (2008). The stem cell niche. Journal of Pathology 217, 169–180.

52. Warder, B. N., Nelson, K. A., Sui, J., Anllo, L. and DiNardo, S. (2024). An actomyosin network organizes niche morphology and responds to feedback from recruited stem cells.

53. Wawersik, M., Milutinovich, A., Casper, A. L., Matunis, E., Williams, B. and Van Doren, M. (2005). Somatic control of germline sexual development is mediated by the JAK/STAT pathway. Nature 436, 563–567.

54. Wingert, L. and DiNardo, S. (2015). Traffic jam functions in a branched pathway from Notch activation to niche cell fate. Development (Cambridge) 142, 2268–2277.

55. Yamashita, Y., Jones, L. and Fuller, M. (2003). Orientation of Asymmetric StemCell Division by the APC TumorSuppressor and Centrosome. Science (1979) 301, 1547–1550.

56. Yamashita, Y. M., Fuller, M. T. and Jones, D. L. (2005). Signaling in stem cell niches: Lessons from the Drosophila germline. J Cell Sci 118, 665–672.

57. Zamfirescu, A-M., Yatsenko, AS., and Shcherbata, HR. (2022). Notch signaling sculpts the stem cell niche. Front. Cell Dev. Biol. 10:1027222

58. Zheng, Q., Wang, Y., Vargas, E. and DiNardo, S. (2011). Magu is required for germline stem cell self-renewal through BMP signaling in the Drosophila testis. Dev Biol 357, 202–210

